# Membrane specificity of the human cholesterol transfer protein STARD4

**DOI:** 10.1101/2023.12.04.569895

**Authors:** Reza Talandashti, Larissa van Ek, Charlotte Gehin, Dandan Xue, Mahmoud Moqadam, Anne-Claude Gavin, Nathalie Reuter

## Abstract

STARD4 regulates cholesterol homeostasis by transferring cholesterol between plasma membrane and endoplasmic reticulum. The STARD4 structure features a helix-grip fold surrounding a large hydrophobic cavity holding the sterol. Its access is controlled by a gate formed by two flexible loops - Ω1 and Ω4- and the C-terminal α-helix. Besides this, little is known about the mechanisms by which STARD4 binds to membranes and extract/releases cholesterol. All available structures of STARD4 are without a bound sterol and display the same closed conformation of the gate. The cholesterol transfer activity of the mouse STARD4 is enhanced in the presence of anionic lipids, and in particular of phosphatidylinositol biphosphates (PIP2) for which two binding sites were proposed on the mouse STARD4 surface. Yet only one of these sites is conserved in human STARD4. We here report the results of a liposome microarray-based assay and microseconds-long molecular dynamics simulations of apo-and holo forms of human STARD4 with complex lipid bilayers mimicking the composition of the donor and acceptor membranes. We show that the binding of apo form of human STARD4 is sensitive to the presence of PIP2 through two specific binding sites, one of which was not identified on mouse STARD4. We report two novel conformations of the gate in holo-STARD4: a yet-unobserved close conformation and an open conformation of Ω4 shedding light on the opening/closure mechanism needed for cholesterol uptake/release. Overall, the modulation of human STARD4 membrane-binding by lipid composition, and by the presence of the cargo supports the capacity of human STARD4 to achieve directed transfer between specific organelle membranes.

## Introduction

Cholesterol plays a critical role in many cellular signaling processes, and it is necessary for cells to maintain specific levels of cholesterol in each subcellular membrane(1). However, vesicular transport is an inefficient method for distributing lipids, including cholesterol, to subcellular membranes(2, 3). Moreover, due to its entropic nature, cholesterol cannot diffuse through the cytosol to reach its target membrane(3, 4).

Lipid transfer proteins (LTPs) transfer lipids between intracellular membranes through non-vesicular mechanisms, enabling rapid changes in lipid levels within membranes(3, 5). One major group of LTPs in human is the steroidogenic acute regulatory protein (StAR)-related lipid-transfer (StART) domain, which consists of 15 domains categorized into six groups(3, 6, 7). Among these, StAR-related lipid transfer domain protein 4 (STARD4) is a soluble single domain protein that facilitates cholesterol trafficking between the plasma membrane, endocytic recycling compartment, and endoplasmic reticulum(8–13). It has been suggested to be an important player in regulating cholesterol homeostasis in human cells(12). Unlike many other LTPs, STARD4 does not have a membrane-targeting domain (e.g. PH domains), hence the activity of STARD4 is not limited to the membrane contact sites (MCSs)(7). The propensity of STARD4 to bind to membranes and its cholesterol transfer function have been shown to depend on membrane composition. In particular increasing the degree of unsaturation of the lipid acyl chains increases sterol transfer activity of mouse STARD4(12). Furthermore the presence of negative lipids such as phosphatidylserine (PS), phosphatidylinositol (PI) and phosphatidylinositol biphosphates (PIP2; PI(4,5)P2 and PI(3,5)P_2_) in both donor and acceptor liposomes increases the rate of transfer of dihydroxyergosterol in a FRET assay, compared to neutral liposomes(12–14).

STARD4 shares with the other START domains the helix-grip fold characterized by 9 beta strands forming a sheet around the hydrophobic cavity that holds the lipid cargo (Fig S1A & S1B). The cavity is further closed by the 20 amino acids-long C-terminal helix (C-ter helix), and the so-called Ω1 loop (β5-β6) and Ω4 loop (β9-C-ter helix). Conserved residues within the cavity are responsible for lipid cargo specificity within the STARD family of LTPs(15). The Ω1 loop is thought to function as a lid and is necessary for cholesterol transfer activity. No structure of STARD4 (or other eukaryote sterol-transferring START domain) with a bound sterol is available but a few models have been reported showing that one sterol molecule can fit in the cavity with its hydroxyl group directed towards the protein core(14, 16, 17). In all available structures of mouse or human STARD4(10, 11, 13) the C-ter helix, Ω1, and Ω4 loops are in similar conformations and defined as a closed state since the hydrophobic pocket is not accessible to cholesterol. Because of the lack of diversity in the experimentally determined structural ensemble of STARD4, little remains known about the conformational changes undergone by the domain. However, in structures of other STAR-related sterol transfer domains of the yeast LAM4(18) and Ysp2(19) proteins the Ω1 loop appears in different conformations. In particular the tip of the Ω1 loop shifts towards the C-terminal helix in both holo forms shielding the cargo from the solvent. Moreover, NMR 15N spin-relaxation and amide exchange along with molecular dynamics simulation of the human STARD6 protein revealed concerted fluctuations of the Ω1 loop and C-terminal helix allowing the opening of the binding pocket(20). These data provide grounds to expect that the STARD4 Ω1 loop also might be a flexible lid.

**Figure 1.**
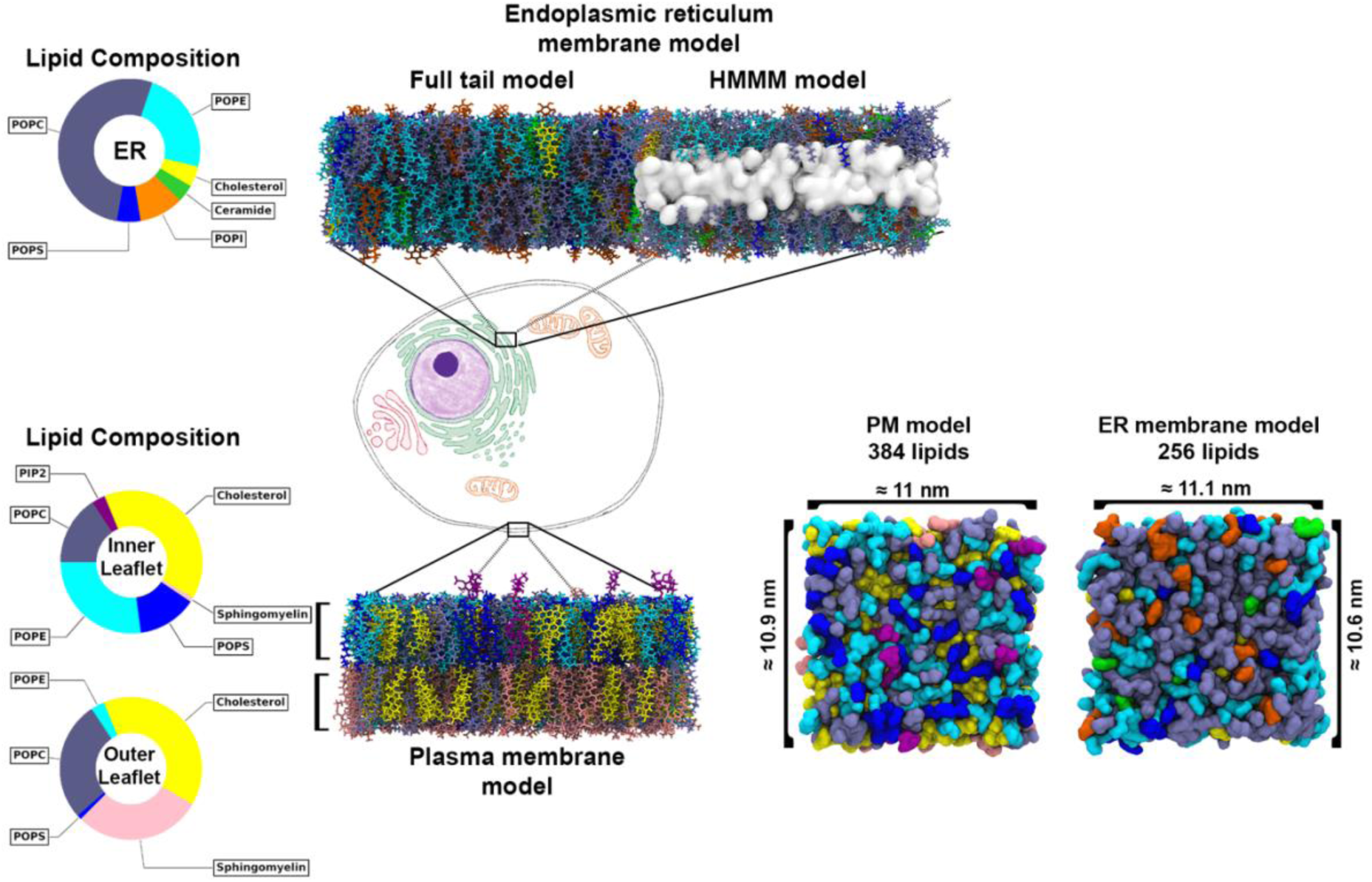
The lipid bilayer models used in this study and their lipid composition.

Sterol transfer and membrane binding assays of the wild type protein and mutants have shed light on several aspects of the structure-activity relationship of STARD4, using either the mouse STARD4 or the human protein which share 79% sequence identity overall. Most substitutions are in loop regions with notably only 1 substitution (W/L) in Ω1 and none in Ω4 (Fig S1C). In particular the importance of the Ω1 loop for transfer of sterol between liposomes has been shown in both mouse STARD4 and human STARD4; mutations in Ω1 either abolish(11) or reduce(13) STARD4 transfer activity in liposomes, depending on the position and nature of the substitution. The deletion of 4 amino acids (of which 3 are conserved) in Ω1 did abolish sterol transfer in the human protein(11). Results from a fluorescence-based membrane penetration assay indicated that the Ω1 loop in mouse STARD4 is not involved in membrane binding. Indeed the Ω1 loop did not penetrate between lipid head groups in liposomes containing phosphatidylserine (PS), phosphatidylcholine (PC), phosphatidylethanolamine (PE) and cholesterol(13), whereas the C-ter helix does penetrate in this assay. Yet, the sequence of Ω1 in human STARD4 is characteristic of a membrane-binding segment(21) with three hydrophobic residues (Leu107, IIe110, Ile111) and one tryptophan (Trp108, Fig S1), which is a typical interface residue(21, 22). The sequence of mouse STARD4 Ω1 is almost identical to that in human STARD4 yet with the notable exception of Trp108 being substituted by a Leu in the mouse protein (Fig S1C).

The increased mouse STARD4 membrane affinity and transfer activity observed with PIP2-containing liposomes are likely to be explained by the presence of two PIP2 neighboring binding sites located on the β1 and β2 strands (Arg46, Lys49, Lys52, Arg58) and at the tail end of the C-terminal helix (Ser215, Arg218, Arg219, Arg222). These sites were identified using mutagenesis and molecular dynamics simulations(14), in a recent study which also suggests that the Ω1 loop might be involved in membrane binding unlike what was suggested by Iaea *et al*(13). Interestingly the first site is conserved between mouse and human STARD4 and corresponds to residues Arg30, Lys33, Lys36, and Arg42 (Fig.S1C). In the second site only Arg218 and Lys219 are conserved between mouse and human where the equivalent amino acids in human STARD4 are Arg202 and Lys203. Ser215 in mouse STARD4 is replaced by a glycine (Gly199) and Lys219 has no equivalent in human STARD4 (Fig S1C), making it unlikely that human STARD4 has a PIP2-binding site equivalent to that of mouse STARD4 on the C-terminal helix.

To evaluate the membrane specificity of human STARD4, we here present the results of liposome microarray-based assay (LiMA) and microseconds-long molecular dynamics simulations of the human STARD4. Molecular dynamics simulations have proven useful to provide reliable models of interfacial protein-lipid recognition(23–26) and conformational changes(27, 28). The simulations included human STARD4 -either in its apo or cholesterol-bound form-in the presence of either water, or complex lipid bilayers. The bilayer compositions were chosen to mimic the composition of the plasma membrane and of the endoplasmic reticulum membrane. This allowed us to map protein-lipid interactions and increase our understanding of the lipid specificity of the human STARD4 and the impact of the sequence difference with the mouse protein on PIP2 dependency. The simulations also allow us to probe conformational changes of the Ω1 loop and of the gate entrance region, as well as the role of the Ω1 loop in membrane binding. Finally, we could also gain insights into the allosteric role of the lipid bilayers on the gate opening.

## Results

### 1. Simulations of apo and holo STARD4 in water

As there is no experimental structure of STARD4 with cholesterol bound, we used molecular docking to build a model structure for the STARD4-cholesterol complex, subjected it to MD simulations and compared the results to simulations of the apo form.

During the simulations, the STARD4 backbone deviates from the X-ray structure by less than 2.2 Å (Cf plot of root mean square deviation (RMSD) on Fig 2A) and this is irrespective of the presence of cholesterol in the lipid binding site. Cholesterol docked at the far end of the binding pocket opposite to the entrance gate located between the Ω1 and Ω4 loops. The pocket mostly consists of nonpolar side chains and a few polar residues located at the gate and bottom of the cavity (Fig 2B). We measured the distance between the cholesterol hydroxyl group and the Ser120 sidechain of STARD4 to estimate cholesterol movement in the binding pocket (Fig 2D). The cholesterol hydroxyl group, which originally forms hydrogen bonds with Ser120 and Ser131 (Fig 2C), then moves towards the gate after about 250 ns in one replica and 1.3 μs in the second replica simulations (Fig 2D). By the end of the simulation, the tail of cholesterol was positioned between the Ω1 and Ω4 loops within a cluster of hydrophobic amino acids consisting of Ile111 (Ω1), Leu177(Ω4), Val186(Ω4) and Ile181(Ω4), and also in contact with Phe116 on β5. The cholesterol head retained some flexibility through conical movements. It made occasional hydrogen bonds with the sidechain of Arg76, Trp155, and the backbone of Ile70 (Fig 2C), similar to recently reported simulations of mouse STARD4 in water(17). Visual inspection of the MD trajectories of holo STARD4 revealed a conformational change of the Ω1 loop (Fig 3) whereby residues 104-110 of Ω1 are displaced towards the C-terminal helix. This change can be characterised using the radius of gyration (Rg) of Ω1 and the number of contacts between Ω1 and the C-terminal helix, as shown by the scatter plots on Fig 3A and Fig 3B for the apo and holo forms, respectively. We observe only one state for the apo form of the protein, with an Rg value of around 5 Å and less than 180 contacts between atoms of the Ω1 loop and atoms of the C-terminal helix (Fig 3A). In contrast, the holo structure samples two major states with significant increase in the Rg value and number of loop-helix contacts (Fig 3B). The Rg value of the Ω1 loop along the trajectories displayed an increase in the two holo replica simulations (Fig 3C). These findings suggest that the Ω1 loop undergoes conformational expansion in the holo structure (Fig 3D), which could serve as a gate-closing mechanism to protect cholesterol from the aqueous environment.

**Figure 2.**
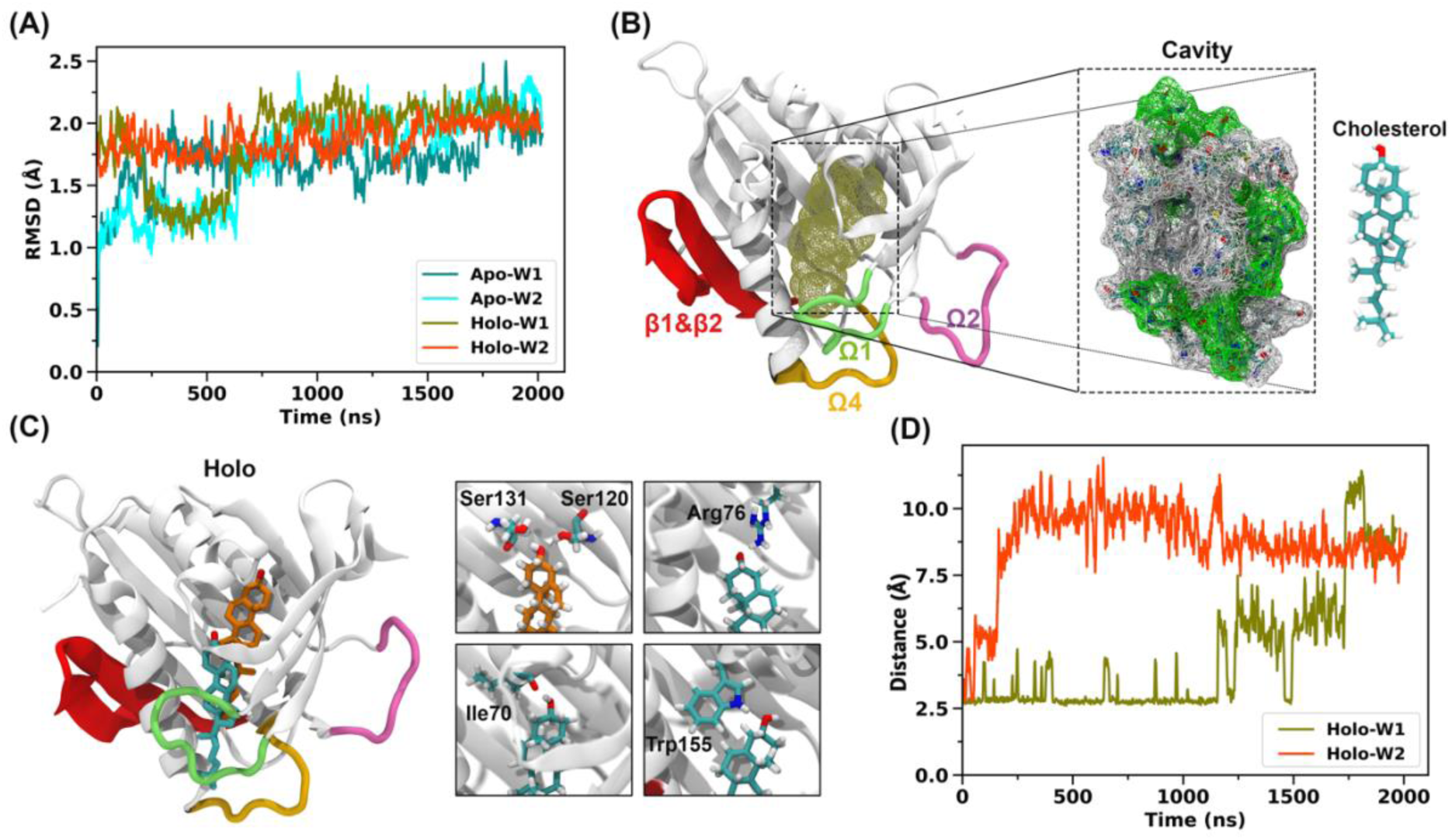
Position of cholesterol in STARD4. **(A)** sProtein backbone RMSD in the apo and holo states relative to the X-ray structure. **(B)** Structure of cholesterol (tan-colored mesh) docked into apo STARD4 with β1-β2 (red), Ω1 (lime), Ω2 (mauve) and Ω4 (orange) highlighted. The residues forming the cavity are shown in a mesh model on the right-hand side, using green and gray for polar and nonpolar residues, respectively. **(C)** Structure of STARD4 with cholesterol bound before MD simulation (docking pose, brown) and after MD simulation (blue). **(D)** distance between cholesterol OH group and Ser120 sidechain along the two MD simulation replicas.

**Figure 3.**
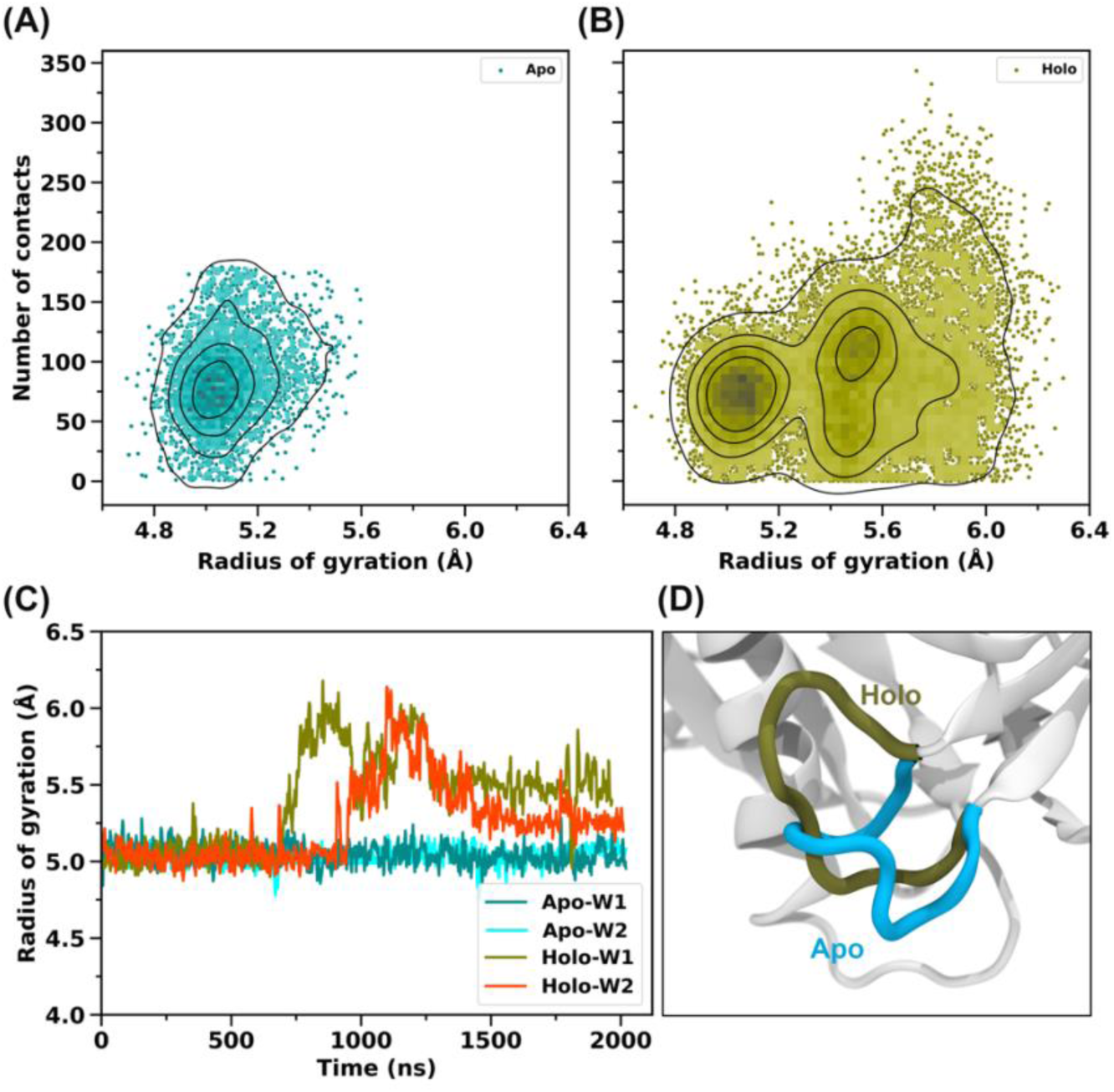
The Ω1 loop closes the gate upon binding of cholesterol. **(A-B)** Scatter plot showing Ω1 radius of gyration (Rg) and number of contacts between Ω1 and the C-terminal helix in the apo and holo simulations. **(C)** Rg of the Ω1 loop over time in simulations. **(D)** Conformational changes in the Ω1 loop of STARD4 in apo and holo forms, highlighting the gate-closing mechanism in the holo form.

Furthermore the first turn of the C-ter helix (Gln183-Val186) as observed in the X-ray structure or in the simulations of apo-STARD4 is unwinded in the simulations of holo form and the amino acids Gln183, Ser184, Ala185 and Val186 are now extending the Ω4 loop (Cf Supp Inf Fig S2).

### 2. Dependence of STARD4 binding orientation on cargo and bilayer composition

#### Membrane-bound forms of apo and holo STARD4 to bilayers mimicking the plasma membrane and endoplasmic reticulum membranes

STARD4 in its apo form extracts cholesterol from the plasma membrane to later deliver it to the endoplasmic reticulum. In order to generate models of the membrane-bound forms of STARD4, we conducted 2 μs-long simulations of apo-STARD4 on the nPM bilayer and of holo-STARD4 on pER, which are models for the plasma membrane and the ER membrane, respectively. The lipid composition of each bilayer is reported in Table 2 (Methods section). Briefly nPM is an asymmetric bilayer consisting of POPC, POPE, POPS, PI(4,5)P2, PSM and cholesterol, where PI(4,5)P2 is only present in the inner leaflet exposed to STARD4. pER is symmetric and contains only POPC, POPE, POPS, POPI, CER180 and cholesterol. We report the RMSD of the protein backbone with respect to the X-ray structure (Fig S3) and the minimum protein-lipid distance (Fig S4). The apo and holo protein structures do not deviate much from the X-ray structure (2.4 Å at most) though the RMSD is slightly higher than for the simulations of STARD4 in water. Both forms bind to the bilayer within at most 200 ns (Fig S4). The apo and holo forms bind to the nPM and pER bilayers in a very similar fashion, engaging four regions (β1&β2, Ω1, Ω2, and Ω4) in interactions with the membrane (Fig 4A). Of these, β1&β2, Ω1, and Ω2 mainly interact with the headgroup and phosphate regions of the lipids without significantly penetrating the lipid bilayers. The loop Ω4 however penetrates deeply into the lipid bilayer positioning the gate of the cavity (the region between Ω1 and Ω4 loops) at the membrane interface (Fig 4A & 4B).

**Figure 4.**
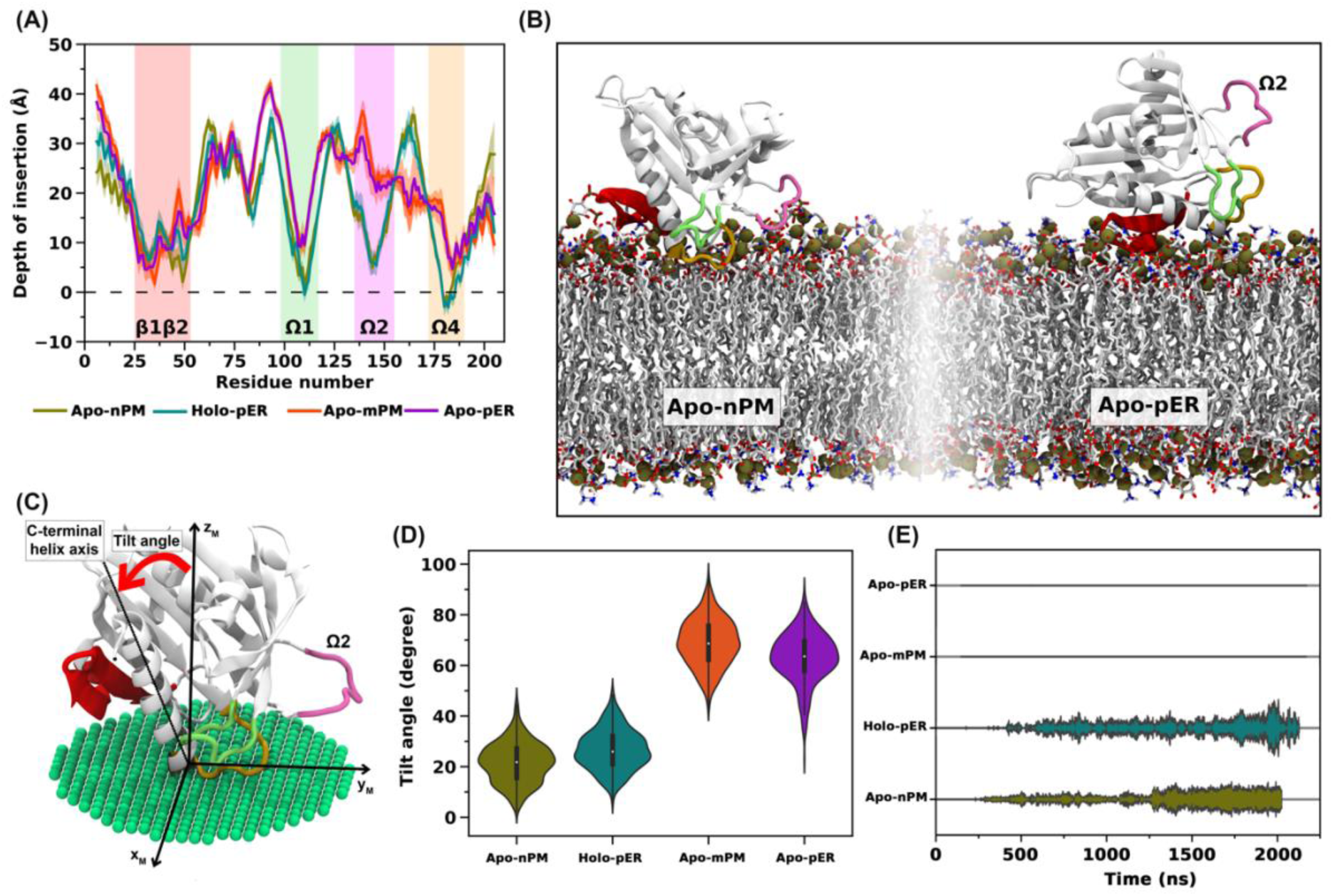
Membrane binding modes of STARD4 in Apo-nPM, Holo-pER, Apo-mPM and Apo-pER simulations. The composition of each lipid bilayer is given in Table 2 (Methods section**) (A)** depth of insertion of amino acids along the STARD4 amino acids sequence in four systems. The shaded area represents the standard deviation of data. The data are averaged over the last 500 ns of all replicas. **(B)** Last snapshot of the bound apo STARD4 on the nPM (left) and pER (right) bilayers. The protein and lipids are represented using cartoon and licorice models, respectively. The carbon, oxygen, and nitrogen atoms are colored white, red, and blue, respectively, and the phosphorus atoms of lipids are shown as tan-colored spheres. **(C)** Definition of the protein tilt angle. **(D)** Tilt angle and **(E)** the kite diagram of number of contacts between Ω2 loop and bilayers along the four simulations.

#### Orientation of apo STARD4 and lipid bilayer composition

To investigate the dependence of STARD4 on the presence of PI(4,5)P2 lipids, we simulated the apo form in the presence of a plasma membrane-like bilayer stripped from its PIP2 lipids (Cf mPM in Table 2). Without PIP2 lipids the protein adopts a strikingly different orientation (Fig 4A) and a more superficial anchoring than in the presence of PI(4,5)P2 (Fig 4B). The Ω2 loop remains above the bilayer and the gate to the hydrophobic cavity faces the aqueous solution instead of being at the bilayer interface. The tilt angle -defined as the angle between the normal to the membrane plane (z-axis) and the C-terminal helix axis (Fig 4C) – is low in the presence of PI(4,5)P2 (around 20 degrees). In the absence of PI(4,5)P2 the C-terminal helix is nearly parallel to the membrane surface and the tilt angle is higher (60 to 80 degrees) (Fig 4D). While the Ω2 loop of apo STARD4 maintains constant contact with nPM lipid bilayer model, these are systematically absent in the absence of PIP(4,5)P_2_ lipids (Fig 4E). The same orientation is observed when apo STARD4 is simulated in the presence of a simple POPC bilayer (Fig S5D) or with the ER-like membrane (pER): the tilt angle is high and no contacts between Ω2 and lipids are observed (Fig.4A-B and D-E). In summary we observe two distinct binding modes of the apo STARD4 where only one seems conducive to cholesterol uptake with the gate facing the bilayer surface. This orientation requires the presence of PI(4,5)P2.

#### Orientation of holo STARD4 and bilayer composition

The holo form of STARD4 appears less sensitive to membrane composition than the apo form. Simulations of holo STARD4 on the simple ER-like bilayer (pER) led to the same orientation (Fig 4A&4D) as that of the apo form on nPM where Ω2 contacts the lipids (Fig 4E); the gate is positioned at the membrane interface. Interestingly, the same orientation is obtained for the holo form on the nPM bilayer (Fig S5A).

Since STARD4 has been shown to have higher cholesterol transfer activity between liposomes enriched with unsaturated lipids(12), we also simulated STARD4 in the presence of a more fluid bilayer. We used a bilayer similar to pER in its composition but where the 1-palmitoyl-2-oleoyl tails were replaced with 1,2-dioleoyl (except for POPI) (dER in Table1). The resulting binding mode was similar to that observed on the pER bilayer (Cf. Fig S5B). This orientation was also observed in simulations with a complex ER membrane model containing a greater diversity of headgroups and tail types (cER, see Table 2 and Fig S5C). Overall, these results indicate that the presence of cholesterol in the hydrophobic pocket reduces the membrane selectivity of STARD4 at least compared to the apo STARD4.

### 3. Lipid binding sites on STARD4 surface

#### PI(4,5)P2 binding sites in β1β2 and Ω2-Ω4

Upon binding to PIP2-containing bilayers, the STARD4 shows two binding sites for PI(4,5)P2, a primary binding site on the β1 and β2 strands and the secondary binding site in the region between the Ω2 and Ω4 loops (Fig.5A). Both PI(4,5)P2 binding sites are composed of basic residues, with Arg30, Lys33, Lys34, and Arg42 in the primary binding site and Lys141 and Arg178 in the secondary binding site (Fig 5A). Along the simulations STARD4 always establishes hydrogen bonds first with the PIP2 primary binding site, and then with the secondary binding site (Movie 1 and Fig S6). The same PI(4,5)P2 lipids remain hydrogen bonded to the binding sites during the whole trajectories (Fig S6), indicating the specificity of these region towards PIP2 lipids. Arginine residues in both PIP2 binding sites make hydrogen bonds with high occupancy (more than 75%), while the lysine residues have occupancies between 40 and 50% (Fig 5B). These hydrogen bonds mostly occur between the side chain of basic residues and phosphate groups of the inositol ring in the PI(4,5)P2 lipid.

**Figure 5.**
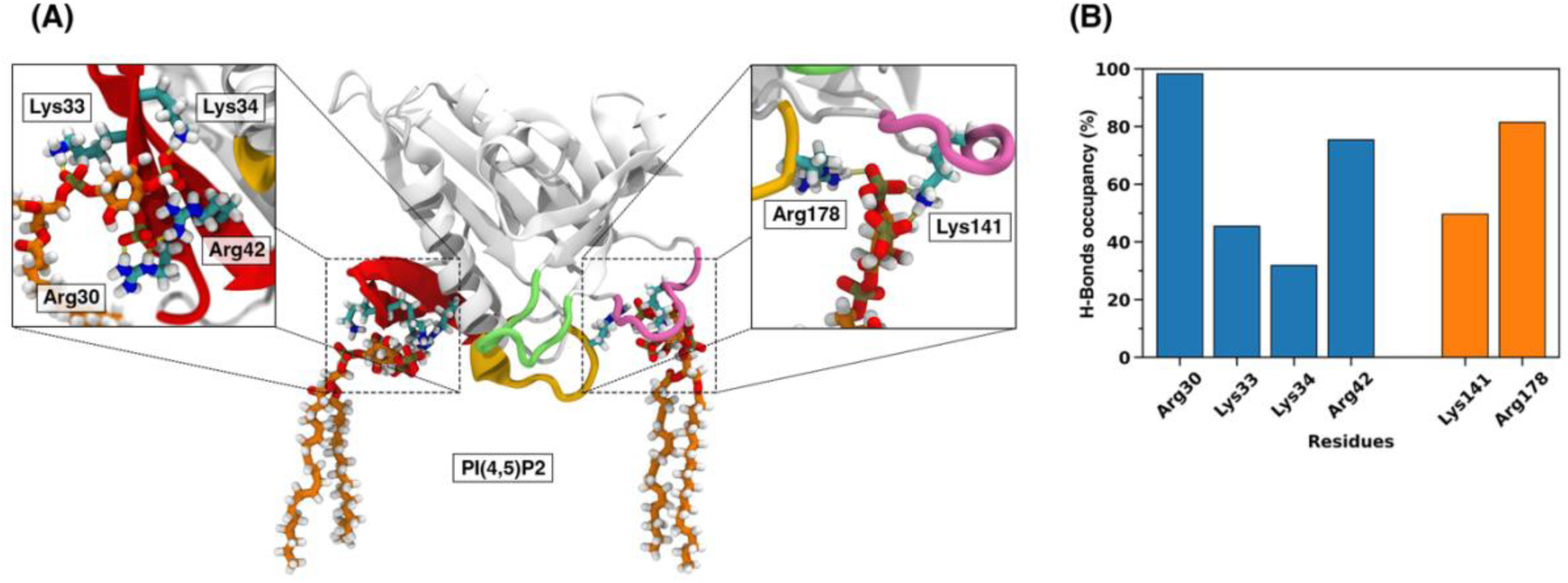
Specific PI(4,5)P2 binding sites. **(A)** location of the STARD4 PI(4,5)P2 binding sites and residues involved. **(B)** occupancies of each hydrogen bond averaged during the last 500 ns of the two replica trajectories.

#### Lipid-protein interactions through the Ω1 and Ω4 loops

Ser112 (Ω1) and Arg114 (β6) engage in long-lasting hydrogen bonds with the bilayer lipids, mostly with phosphate groups (Fig 6A). No particular lipid type seems to be preferred at this site as we observe different types of lipids such as PC, PE, and PS, and there can be exchange of lipids during a simulation. These hydrogen bonds only exist in the binding orientation where the C-terminal helix is almost perpendicular to the membrane surface. Arg114 has hydrogen bond occupancy ranging from 50 to 90% which is higher than the occupancy of Ser112 in all simulation systems. The unsaturated tail of the phospholipid in this position frequently snorkels and tries to insert in the gap between the Ω1 and Ω4 loops (Fig S7). In the orientations with a “vertical” C-terminal helix, the Ω4 loop engages in more hydrogen bonds with the lipids than Ω1 (Table 1). In the holo forms, Ser184 engages in long-lasting H-Bonds with lipids which it cannot do in the apo form where the Ser184 still folded in the C-terminal helix. Visual inspection of our simulations indicates that the unwinded turn of C-ter helix allows for a deeper anchoring of that region than in the apo forms.

**Figure 6.**
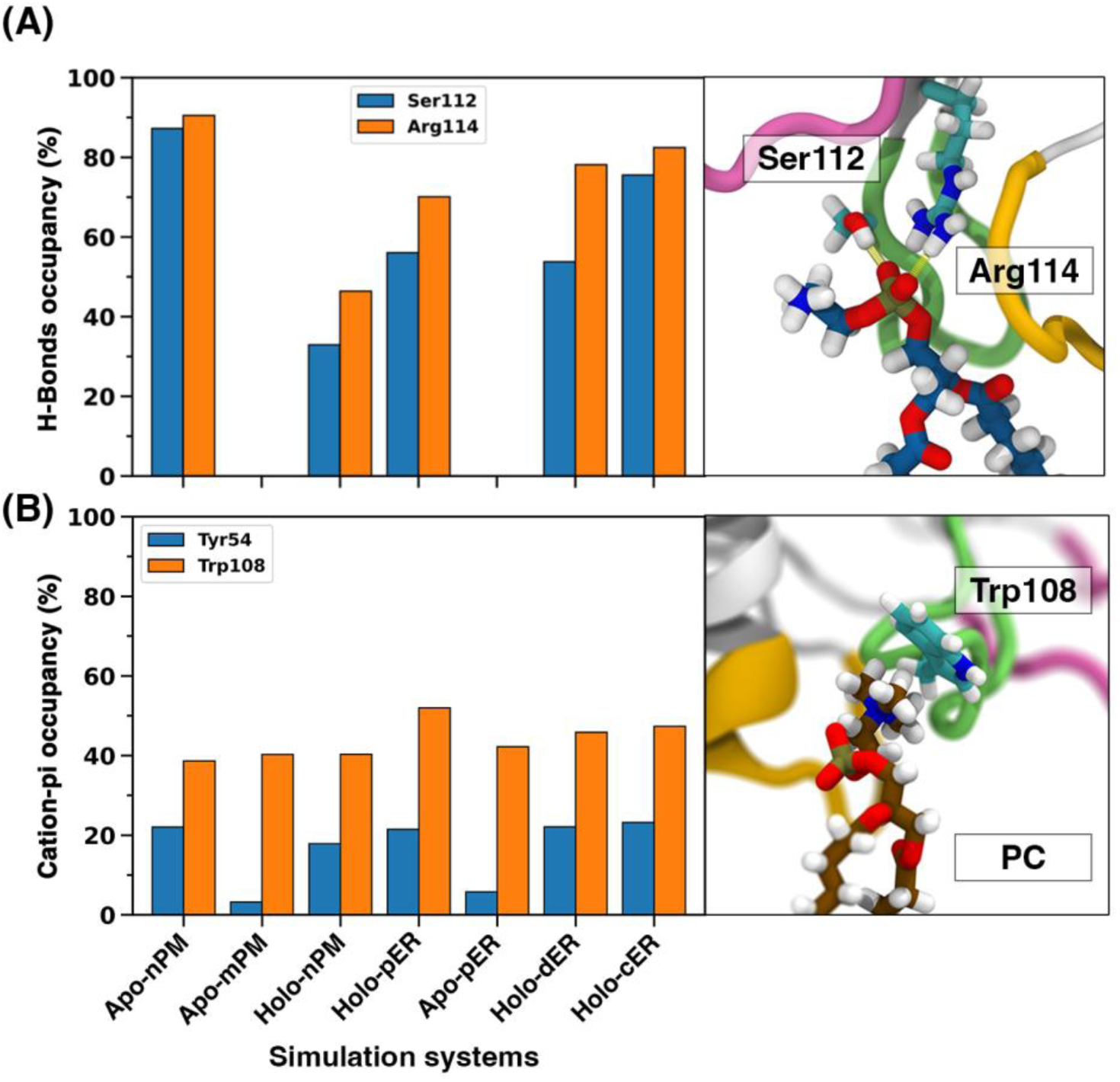
Specific lipid binding sites on the Ω1 loop. **(A)** Occupancy of hydrogen bonds between Ser112 (blue) and Arg114 (orange) and the phosphate group of any lipid in different simulation systems. The values are averaged over the last 500 ns of the two replicates. **(B)** Occupancy of cation-π interactions between any phosphatidylcholine headgroup and Trp108 and Tyr54 residues. The values are averaged over the last 500 ns of all replica simulations.

**Table 1:** Inventory of hydrogen bonds and hydrophobic contacts between bilayer lipids and the Ω1 and Ω4 loops. The occupancy of hydrogen bonds is calculated over the last 500 ns of trajectories and reported if the value is above 25% and present in at least in two replicas. Hydrophobic contacts are averaged number of contacts per trajectory frame over all replicas in each case, calculated during the last 500 ns of trajectories. Average number of hydrophobic contacts per frame are reported if > 0.5.

|  |  | Apo-nPM | Apo-mPM | Holo-<br>nPM | Holo-pER | Apo-pER | Holo-<br>dER | Holo-<br>cER |
| --- | --- | --- | --- | --- | --- | --- | --- | --- |
| Hydrogen bonds (occupancy %) |  |  |  |  |  |  |  |  |
| $\Omega 1$ | Trp108 | - | - | 40.4% | 37.8% | - | 49.7% | 43.7% |
|  | Asn109 | - | - | 27.6% | 42% | - | 29.1% | 35.5% |
|  | Ile110 | - | - | 32.2% | 39.7% | - | 47.1% | 33.4% |
| $\Omega 4$ | Asp176 | 34.8% | - | 28.8% | 34.6% | - | 24.4% | 30.8% |
|  | Arg178 | 61.4% | - | 28.2% | 42.3% | - | 35.2% | 28.7% |
|  | Gly179 | 43.7% | - | 27.1% | 57.7% | - | 38.1% | 47.6% |
|  | Met180 | 50.4% | - | 33.2% | 37.1% | - | 41.9% | 39.5% |
|  | Ile181 | - | - | 36.2% | 32.5% | - | 35.6% | 33.8% |
|  | Gln183 | 35.1% | - | 27.3% | 35.7% | - | 50.7% | 43.2% |
|  | Ser184 | - | - | 82.26 | 56.3% | - | 68.9% | 51.1% |
| Hydrophobic contacts (average contact per frame) |  |  |  |  |  |  |  |  |
| $\Omega 1$ | Ile110 | 0.72 | - | 0.73 | 0.62 | - | 0.75 | 0.68 |
| $\Omega 4$ | Met180 | 2.17 | - | 0.80 | 1.99 | - | 1.14 | 1.57 |
|  | Pro182 | 1.36 | - | 0.81 | 1.72 | - | 1.87 | 1.80 |

Furthermore, we mapped the hydrophobic contacts between STARD4 and the lipid bilayer models (Table 1). The results show that Ile110 in Ω1 loop and Met180 and Pro182 residues in Ω4 loop establish hydrophobic contacts with the lipids tails in those simulations where the C-terminal helix is almost perpendicular to the membrane surface.

Finally, Trp108 (Ω1) also engages in cation-pi interactions with phosphatidylcholine headgroups. The occupancy is not very high (40-50%) but the interaction is consistently present in every simulation, irrespective of the form of the protein (apo or holo) and of the bilayer used in the simulation (Fig 6B). It is worth noting that Tyr54 (β3 strand) engages in short cation-pi interactions as well but with fairly low occupancies.

### 4. Recruitment of STARD4 on the surface of liposomes

The binding of STARD4 to DOPC, POPC, and PM-mimic liposomes with and without 5 or 10% PIPs was examined using a biochemical assay, the liposome microarray-based assay (LiMA) (Fig 7). Different types of PIPs have been used in this experiment, including PI(3)P, PI(4)P, PI(5)P, PI(3,4)P2, PI(3,5)P2, PI(4,5)P2, and PI(3,4,5)P3. Overall, these results show increased binding of STARD4 to liposomes containing PIPs (Fig 7A) which is also known to enhance the rate of sterol transfer(14). By adding 5 mol% PIPs to DOPC, POPC, and PM-mimic liposomes, a higher binding affinity is seen in PM-mimic liposomes. It shows that STARD4 prefers complex membranes over simple membranes (Fig 7B). At 10 mol% PIPs, a preference starts to arise for DOPC liposomes indicating that STARD4 prefers fluid membranes over rigid membranes at high PIPs concentrations (Fig 7C). Overall, STARD4 tends to bind PM mimics with 5 mol% PIP, PIP2 and PIP3, and DOPC membranes with 10 mol% PIPs. All other conditions were distributed around the log(FC) value of 0.

**Figure 7.**
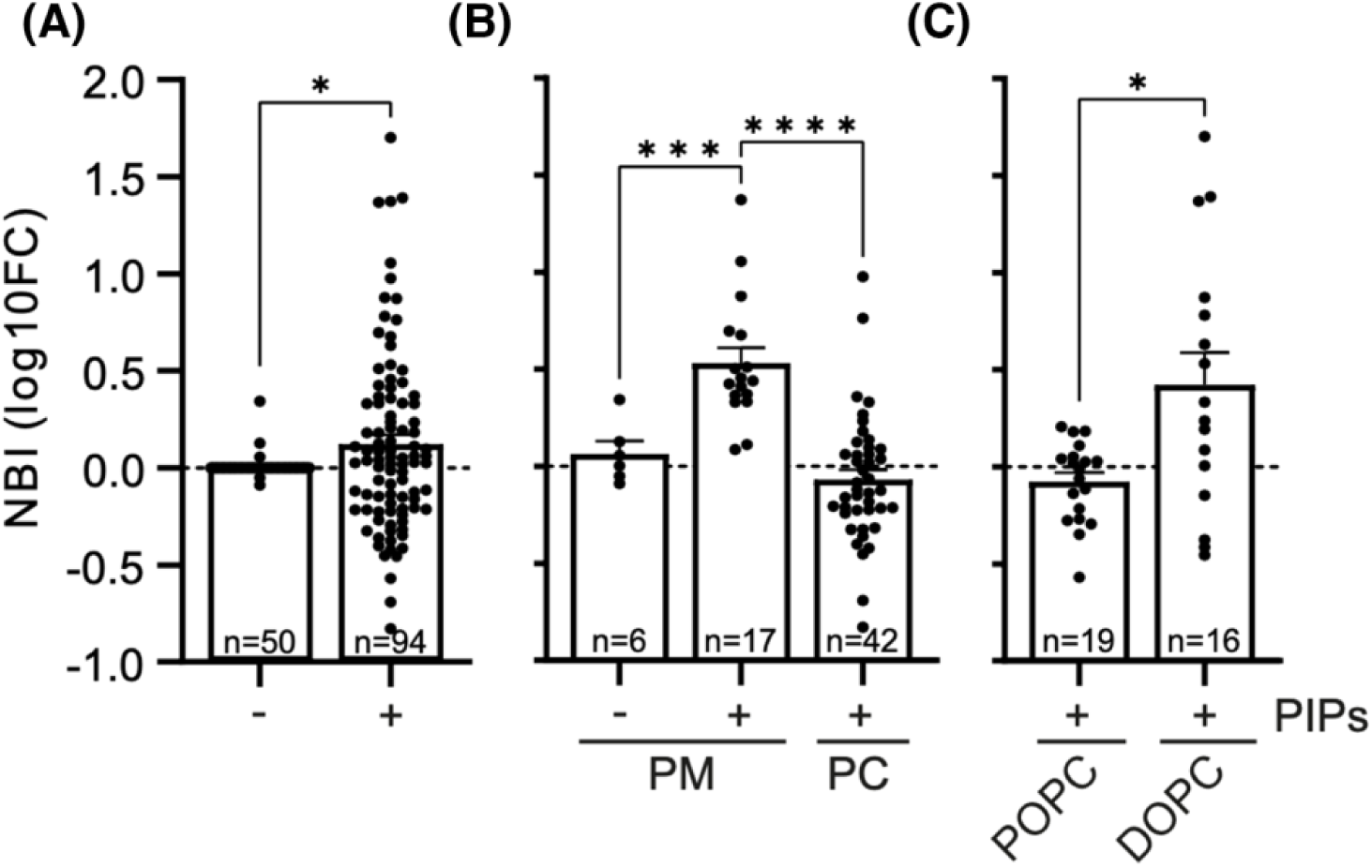
STARD4 binds PIPs in general. **(A)** Comparison of cumulative DOPC, POPC, PM mimic membranes vs DOPC, POPC, PM mimic membranes with 5 or 10 mol% PIPs (PIP1, PIP2, PIP3 conditions all included). **(B)** Comparison of plasma membrane mimic with or without 5 mol% PIPs (PIP1, PIP2, PIP3 conditions all included) and PC (DOPC or POPC) with 5 mol% PIPs. **(C)** Comparison of 10 mol% PIPs (PIP1, PIP2, PIP3 conditions all included) in a DOPC or POPC membrane. All the NBI values are expressed as the 10log(fold change) with its corresponding PC (DOPC or POPC) to account for chip-to-chip variability. Either an unpaired t-test with Welch’s correction or a Brown-Forsythe ANOVA test with subsequent Dunnett’s t-test for multiple comparisons was applied. * P<0.05; ** P< 0.01; *** P<0.001; **** P<0.0001. NBI = normalized binding intensity; PM = plasma membrane; PIPs = phosphoinositides; PC = phosphatidylcholine; DOPC = dioleoyl-phosphatidylcholine; POPC = dipalmitoyl-phosphatidylcholine.

### 5. Gate opens at membrane interface

Numerous studies have emphasized the occurrence of local dynamic and structural alterations in peripheral membrane proteins triggered by their interaction with the membrane(28–31). The flexibility of the membrane binding site, in particular the gate region in START domains has been found to be crucial for lipid transfer activity(3). Upon a more detailed examination of the gate region, specifically focusing on the Ω1 and Ω4 loops, in the protein-membrane simulations, we noticed that the gate region shows more flexibility in the holo form compared to the apo form due to the conformational change of the first turn of the C-terminal helix in the holo structure (Fig S2). To quantify this flexibility we report the variation of distance between the Cα atoms of Ile110 (Ω1) and Pro182 (Ω4) in all STARD4-membrane simulations on Figure 8. The results reveal a greater variation in the distance between the Ω1 and Ω4 loops in the holo structure than in the apo (as depicted in Fig 8A), suggesting that the gate is more susceptible to opening in the presence of cholesterol. Among all simulations of holo STARD4 and membrane, protein shows greater variations in the gate region when it is interacting with dER membrane. This observation serves as a driving factor for us to conduct simulations of the holo structure in the presence of a highly mobile membrane mimetic (HMMM) model similar in composition to the pER lipid bilayer and named hER (Cf Table 2). The HMMM model represents lipid bilayers using short tails lipids and a layer of organic solvent to mimick the membrane core, thereby accelerating lipid diffusion and also potentially interfacial processes. Interestingly, we observed a conformational change in Ω4 loop which moves away from Ω1 loop and opens the pocket entrance at the membrane interface (Fig 8B & 8C). Ω1 loop does not show significant conformational change during opening process. The average distance in closed form is about 13Å compared to 19Å in open form. Although cholesterol remains within the cavity, it establishes contacts with bilayer lipids. These contacts are also present in the full-tail membrane simulations. The HMMM simulation indicates that the opening is influenced by the lipids.

**Table 2.** Lipid composition of membrane models used in this study.

| nPM and mPM<sup>1</sup> (384 lipids) |  |  |  |  |  |  |  |  |  |  |  |  |
| --- | --- | --- | --- | --- | --- | --- | --- | --- | --- | --- | --- | --- |
| Lipid type | POPC | POPE | POPS | PI(4,5)P2 | PSM |  |  |  |  |  |  | Chol |
| Outer leaflet | 27% | 4% | 1% | 0% | 39% |  |  |  |  |  |  | 39% |
| Inner leaflet | 16% | 27% | 14%<sup>1</sup> | 3%<sup>1</sup> | 1% |  |  |  |  |  |  | 39% |
| pER membrane model (256 lipids) |  |  |  |  |  |  |  |  |  |  |  |  |
| Lipid type | POPC | POPE | POPS | POPI | CER180 |  |  |  |  |  |  | Chol |
| Each leaflet | 53% | 23% | 5% | 10% | 4% |  |  |  |  |  |  | 5% |
| dER membrane model (256 lipids) |  |  |  |  |  |  |  |  |  |  |  |  |
| Lipid type | DOPC | DOPE | DOPS | POPI | CER180 |  |  |  |  |  |  | Chol |
| Each leaflet | 53% | 23% | 5% | 10% | 4% |  |  |  |  |  |  | 5% |
| cER membrane model (256 lipids) |  |  |  |  |  |  |  |  |  |  |  |  |
| Lipid Type | POPC | PLPC | SAPC | POPE | PSPE | SAPE | SAPI | SLPI | OLPS | PSM | POPA | Chol |
| Each leaflet | 16% | 18% | 27% | 6% | 7% | 7% | 3% | 3% | 3% | 4% | 1% | 5% |

<sup>1</sup> The mPM model does not contain PIP2 lipids, but contains 17% POPS lipids instead of 14%.

**Figure 8.**
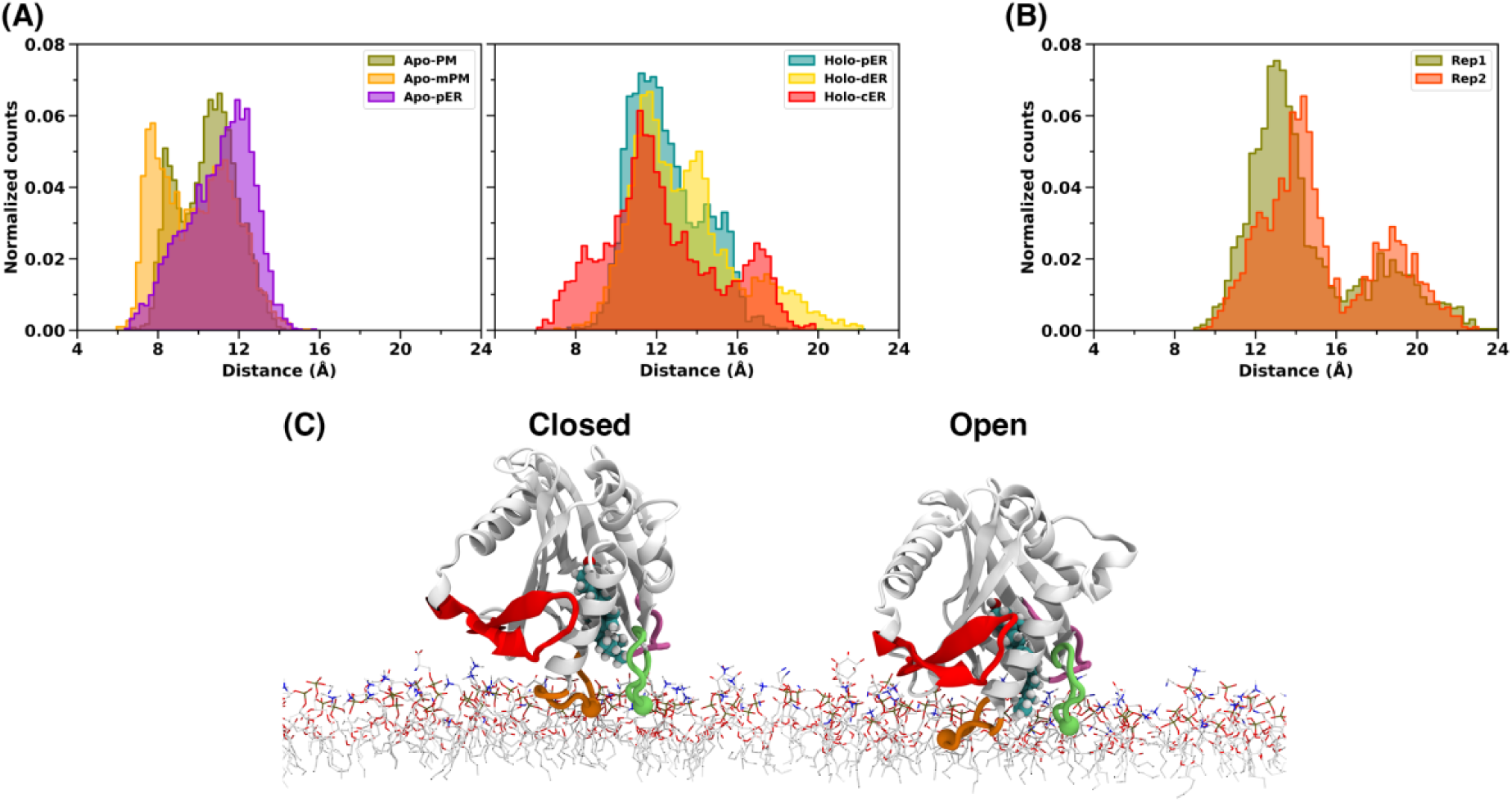
Gate opening at the membrane interface. Distribution of the Ω1-Ω4 distance (Ile110(Cα)-Pro182(Cα) distance) in all STARD4-membrane simulations with **(A)** full-tail membrane models and **(B)** HMMM model (hER model). On the plots the normalized counts are normalized numbers of frames. **(C)** the graphical representation of open and closed state of the protein at the interface of hER lipid bilayer model.

## Discussion

### Structure model of the STARD4-cholesterol complex: unwinding of the C-ter helix and a new closed conformation of the Ω1 loop

The existing X-ray structures of human and mouse STARD4 have been resolved without a ligand in the hydrophobic cavity. We here propose a model for the holo form of human STARD4 using molecular docking and extensive simulations both in an aqueous environment and at the surface of complex lipid bilayers. The position of cholesterol in human STARD4, where the hydroxyl group is hydrogen bonded to Ser120 and Ser131, is similar to the one proposed by Murcia *et al.*(16) for STARD3 where Arg351 and Ser362 correspond to Ser120 and Ser131, respectively. Along the simulations we observe a progressive release of cholesterol from this binding site and towards the cavity gate where the cholesterol aliphatic chain is locked in place within a hydrophobic cluster formed by sidechains of the Ω4 and Ω1 loops. The first turn of the C-ter helix unwinds thereby extending the Ω4 loop. Moreover, the Ω1 loop adopts a closed conformation where amino acids 104-110 are displaced towards the C-terminal helix. This conformation resembles the one of the yeast Ysp2 protein(19). This position of Ω1, consistently observed in simulation replicas, is different from the closed form observed in X-ray structures of the apo form. This conformational change of Ω1 reinforces the idea that Ω1 acts as a lid for the cargo binding site and is important for transfer. Indeed Tan *et al.*(11) showed using dehydroergosterol and a FRET-based transfer assay that the deletion of amino acid 107-110 abolished transfer activity by human STARD4, while the L107R mutation decreased transfer. Iaea et al. used an assay similar to that of *Tan et al*, but on mouse STARD4 and showed that a mutation of the Ω1 residue L124D (equivalent to W108 in human STARD4) abolished transfer activity(13) Using NMR and size-exclusion chromatography they could also show weaker interactions with liposomes which they attribute to a loss of flexibility of mouse STARD4 Ω1 impairing its ability to open.

### Dependence of the membrane-bound orientation on cargo and membrane lipid composition indicates that human STARD4 is able to perform directed cholesterol transfer

We observed two orientations of apo STARD4 on bilayers. One presents the C-ter helix as nearly parallel to the bilayer and the gate to the pocket is not in contact with the bilayer, hence this orientation does not seem conducive to cholesterol uptake. In the second orientation the Ω1 and Ω4 loops which form the gate to the cholesterol binding site are located at the membrane interface. This orientation of human STARD4 requires the presence of PIP2 lipids as was suggested for the mouse STARD4 by Zhang *et al.*(14) who also identified two PIP2 binding sites using molecular simulations. In apo-STARD4 simulations we consistently identify two PI(4,5)P2 binding sites. One is similar to the site identified on mouse STARD4 and consists in conserved amino acids of the β1 (Arg30, Lys33, Lys34) and β2 strands (Arg42). The second site is located in-between Ω2 (Lys141) and Ω4 (Arg178), unlike for the mouse protein where it was reported to be at the tail end of the C-terminal helix and close to the first site. These two sites need to be occupied for apo human STARD4 to adopt a membrane-bound orientation conducive to cholesterol uptake. The simulations of the protein in the absence of PIP2 lead to not only an orientation non-conducive to cholesterol uptake but also a shallow interaction with the bilayer surface (Fig 4A). Interestingly the LiMA experiments indicate that PIP lipids enhance binding of human STARD4 to liposomes of various compositions (Fig 7) as was shown for mouse STARD4(14). Our work suggests that this decreased affinity might be the result of the alternative shallow binding orientation.

For the holo form, we observe only one orientation similar to that of the apo form in the presence of PIP2s and with the Ω1 and Ω4 loops anchored in the bilayer. In other words, in the presence of cholesterol, STARD4 adopts an orientation which is conducive to cargo release, independently of the lipid composition of the bilayer. These findings highlight how the presence of cholesterol within the cavity alters the protein’s responsiveness to the lipid composition of the bilayer. In a related context, a recent investigation by Druzak *et al.* involving STARD2 demonstrates that the presence of cargo in the cavity can influence the binding of STARD2 to the PPARδ protein(32).

The LiMA experiments also bring new insights into the membrane specificity of human STARD4, namely that STARD4 binds preferentially to complex membranes compared to single-component vesicles, and also tends to prefer lipids with double unsaturation (dioleyl) compared to single unsaturation (palmitoyl oleyl). The latter can be seen in the context of the work of Mesmin et al. who showed that the sterol transfer activity of mouse STARD4 increased with the degree of unsaturation of the lipid acyl chains in acceptor liposomes(12). The LiMA data indicates the binding itself is increased with DOPC compared to POPC. The simulations do not show any dependency of the binding on lipid tails or bilayer fluidity, but this is not expected given the range of simulation times which are below experimental time scales. What the simulations suggest is that the degree of packing of the lipids will influence the propensity of the gate to open with PO-containing bilayers having a restricting effect compared to DO tails (for holo form, Fig.8). This would favor release of cholesterol. Altogether these results suggest that both binding and uptake/release are modulated by membrane fluidity.

### hSTARD4-membrane binding: a major role for Ω4 and superficial interactions by Ω1

In orientations conducive to cholesterol release (or uptake), Ω1 plays a role in anchoring STARD4 at the surface of the lipid bilayers. Indeed our simulations show a few but persistent contacts and hydrogen bonds between amino acids of the Ω1 loop and the lipids. Yet, Ω4 anchors deeper than Ω1 and engage in more interactions with the lipids. In the absence of PIP2 lipids and in the presence of the cargo (holo STARD4), the extension of Ω4 by unwinding of a turn of the C-ter helix seems decisive in maintaining the orientation of STARD4 through insertion of Gln183 and Ser184 as anchors in-between phospholipids. Overall, our simulations suggest that Ω4 plays a major role in membrane-binding while Ω1 has superficial interactions, which is partially in disagreement with experimental data(13). Indeed, using a fluorophore coupled to either L124D in Ω1 or M206C (C-ter helix, equivalent to Met190 in human STARD4), Iaea *et al.*(13) observed that Ω1 doesn’t penetrate the lipid bilayers while the C-ter helix does. Yet the experiment was done on liposomes that did not contain PIP2 lipids, so we hypothesize that the observed insertion of the modified M206C reflects the alternative orientation of STARD4. It is also worth noting that studies on related proteins have reported an involvement of Ω1 in membrane binding; notable examples are STARD11(33) as well as the mitochondrial proteins PRELI(34) and the yeast Ups1(35) where the equivalent loops are called Ω loop and α3 loop, respectively.

## Conclusion

To summarize, we reported the results of comprehensive µs-long atomistic molecular dynamics simulations of the apo and holo forms of human STARD4, on lipid bilayers mimicking the plasma membrane and ER membranes. The simulation results were supplemented by liposome microarray-based assays. We propose a structural model for holo-STARD4 and the associated conformational changes of the gate; an alternative close conformation of the Ω1 loop in the presence of cholesterol, and a yet-unobserved open conformation of Ω4 when the protein is bound to a fluid membrane. Overall, we show that human STARD4 membrane-binding is modulated by lipid composition, and by the presence or absence of the cargo, supporting the capacity of human STARD4 to achieve directed transfer between specific organelle membranes. The membrane lipids play a decisive role in the regulation of STARD4 binding which does not rely on additional domains for localization and regulation unlike other LTPs (STARD3/MLN64, STARD11/CERT, RhoGAP-START)(3, 6).

## Methods

### Simulations of STARD4 in water

#### Apo STARD4 system setup

The coordinates of human STARD4 were taken from the RCSB Protein Data Bank (PDB id: 6L1D)(11). The serine at position 75 was replaced by a cysteine using MODELLER (version 10.2)(36) to conform with the reference sequence (UNIPROT ID Q96DR4). All hydrogen atoms were then added to the structure using the VMD psfgen plugin (version 2.0)(37). The pKa values of all ionizable residues were predicted using PROPKA3(38) to be in their standard protonation state. All histidine residues were protonated on their δ nitrogen, following a thorough visual examination of each histidine residue and an analysis of the adjacent amino acids to identify potential hydrogen bond acceptor or donor atoms near the δ and ε nitrogens.

#### Holo STARD4 system setup

Possible binding conformations of cholesterol in STARD4 were sampled using molecular docking with AutoDock Vina(39, 40). The protein structure was extracted from the last frame of the apo STARD4 simulation and a cholesterol molecule was extracted from an equilibrated lipid bilayer (described in the next section). All input files were prepared using AutoDockTools (ADT version 1.5.6)(41). Gasteiger charges were added to the protein and cholesterol. A grid box was created with its center positioned on the side chain of amino acid Cys153. The dimensions of the grid box were set to 66 points in the X dimension, 98 points in the Y dimension, and 68 points in the Z dimension. A grid point spacing of 0.375 Å was utilized for each dimension. The Lamarckian genetic algorithm was used for docking, and ten top poses were generated. The first-ranked pose was selected for the next steps and the resulting cholesteral-STARD4 structure was solvated and subjected to molecular dynamics simulation.

#### Apo and holo STARD4 simulation protocols

The systems were prepared using Charmm-GUI(42, 43) and the simulation protocols were similar for the apo and holo forms. TIP3P water model(44) and sodium ions were used to soak and neutralize the simulated systems (numbers are reported in the Table S1). The simulations were performed with NAMD2.14 package(45) and the Charmm36m force field(46, 47). The simulation systems underwent 10000 steps of energy minimization employing the conjugate gradients (CG) algorithm, followed by equilibration for a duration of 250 ps in the canonical (NVT) ensemble. Position restraints were applied to all atoms of the protein except hydrogens using a harmonic spring potential in the equilibration step. The force applied to the heavy atoms of the backbone was twice as strong as the force exerted on the sidechains. Subsequently, all STARD4 (apo and holo) in water systems were subjected to a 2 μs simulation in the isothermal-isobaric (NPT) ensemble with an integration time step of 2 fs without any positional restraints. Temperature was maintained at 310 K using Langevin dynamics algorithm with the temperature damping coefficient of 1 ps^-1^. Using a Langevin piston with oscillation period of 50 fs, the pressure was set and controlled at 1 atm. Particle mesh Ewald was used to calculate long-range electrostatics interactions(48), and a switching function was used for the coulombic and Lennard-Jones potentials with a cutoff distance of 12 Å and a switching distance of 10 Å.

#### Simulations of STARD4 on membrane models

STARD4 was simulated on bilayers mimicking the endoplasmic reticulum membrane and the plasma membrane (PM) (Fig 1). In this study we used full-tail membrane models and also the highly mobile membrane-mimetic (HMMM) model. All simulations performed are listed in Table S1.

For each protein-bilayer system 2 independent replicates were simulated except for the Holo-pER where 4 replicates were simulated. The replicates differ by the initial position and orientation of the protein with respect to the lipid bilayer, and by the initial distribution of lipids since the bilayers were generated for each replica by random distribution of the lipids.

#### Preparation of lipid bilayers

The lipid compositions of all lipid bilayer models are listed in Table 2. Three different lipid compositions were used to model the ER membrane(49, 50). Two of the models contain only six lipid types (pER and dER: PC, PE, PS, PI, Cer, Chol) but differ by the number of unsaturated chains (only 16:0-18:1 in pER, while dER also included 18:1-18:1 chains). The third endoplasmic reticulum membrane model (cER) is a more complex mixture also including sphingomyelin and phosphatidic acid lipids, as well as a larger diversity of chains than pER and dER (Table 2). The PM models were designed based on the work from Lorent *et al*(51). We built two models where one contains six phosphatidylinositol biphosphate PI(4,5)P2 lipids in the cytoplasmic leaflet and the second lacks PI(4,5)P2. This was done in order to examine the role of PI(4,5)P2.

The ER membrane models are symmetric lipid bilayers and contain 256 lipids, whereas the PM model is an asymmetric membrane model and contains 384 lipids. Due to the high cholesterol level in the PM models, the lipid bilayer contracts in the XY plane; hence a higher number of lipids were used to build that lipid bilayer. The resulting surface areas are 120 nm^2^ and 117 nm^2^ for the PM and ER membrane models, respectively (Fig 1). The bilayer builder module of CHARMM-GUI was used to create topology and structure files of lipid bilayer models(42, 52). First, both ER membrane models and PM models were simulated using the protocol suggested by Sunhwan Jo et al(53). Briefly, the system is optimized using CG followed by multiple equilibration steps in the NVT and then NPT ensembles during which positional restraints are applied and gradually decreased. Finally a production run is performed in the NPT ensemble during 200 ns. The area per lipid (APL) for POPC in the pER bilayer is 62.3 Å^2^ which is in reasonable agreement with previously reported simulation and experimental data(54–56) for pure POPC bilayers (60.4 ± 3.6 Å^2^). This value for DOPC in dER is slightly higher, 67.6 Å^2^, which is in agreement with previous experimental data(56). Due to the high cholesterol concentration (39%) in the nPM model, the APL for POPC expectedly decreases (51.3 Å^2^). We also prepared a pER bilayer using the highly mobile membrane-mimetic (HMMM) model(57). The HMMM builder module of the CHARMM-GUI was used to create the ER-HMMM model (hER) (43, 58).

The lipid tails (except for cholesterol) were trimmed at the position of acyl carbon number 6 and the space was filled with 1,1-dicholroethane (DCLE). The HMMM model was simulated using the protocol suggested by Yifei Qi et al(58). The hER simulation system was equilibrated under six consecutive NPAT simulation with several restraints applied to lipid tails, DCLE molecules and water. To restrict the fluctuation of short-tailed lipids along the membrane normal and prevent their occasional diffusion into the bulk solution, a weak harmonic potential was applied to the z-positions of C21 and C31 (carbonyl carbon) atoms of HMMM lipids. These restraints were maintained even during the production phase of the simulation as in references(58, 59).

#### Simulations of STARD4 on bilayers

The obtained conformations of STARD4 (apo and holo) were placed above the equilibrated membrane models with the Ω1 loop oriented toward the membrane such that the minimum protein-lipid distance is 1.5 nm. All protein-lipid bilayer systems were simulated with 3.5 nm water height on top and bottom of the system for at least 2μs. All simulations were performed with the NAMD2.14 package(45) and the charmm36m force field(46, 47) plus its charmm-WYF extension for cation-π interactions(60, 61). TIP3P water model(44) and the sodium ions were used to soak and neutralize the simulated systems (Cf Table S1). The systems were subjected to energy minimization for 10000 steps using conjugate gradients. Protein-membrane simulation systems were equilibrated under two consecutive NVT simulations for 250 ps each, with an integration time step of 1 fs and velocity reassignment every 0.5 ps, followed by four NPT simulations of 500 ps each with an integration step of 2 fs. To ensure a gradual equilibration of simulation systems, several restraints were employed during the equilibration process. Harmonic restraints were applied to the ions and all atoms of the protein except for hydrogen, repulsive planar restraints were utilized to hinder water molecules from entering the hydrophobic region of the membrane, and planar restraints were imposed to maintain the position of the membrane’s head groups along the Z-axis. Throughout the equilibration, these restraint forces were progressively reduced, allowing the system to achieve a more stable equilibrium state. After equilibration, all systems were simulated in the NPT ensemble without restraint with an integration step of 2 fs. Temperature was maintained at 310 K using Langevin dynamics with the temperature damping coefficient of 1 ps^-1^. Using a Langevin piston with oscillation period of 50 fs, the pressure was set and controlled at 1 atm. All bonds were constrained using the SHAKE algorithm and particle mesh Ewald was used to calculate the long-range electrostatic interactions(48). The charmm36m NB-fix corrections were used for ions. The van der Waals (vdW) interactions were calculated using Lennard-Jones switching function with cutoff of 12 Å and switching distance of 10 Å. The cutoff for electrostatics interactions was set to 12 Å.

#### Trajectory analysis

All analyses were done using MDAnalysis package (version 2.4.2)(62, 63), except for secondary structure analysis and interactions analyses. The secondary structure analysis is done by DSSP package (version 3.0.0)(64, 65). To calculate hydrophobic contacts, hydrogen bonds and cation-pi interactions between protein and lipids we developed a parallel python3 code. The code reads DCD trajectories files and can be run on modern multi-core computers. The code is based on MDAnalysis(50, 51) for parsing of structure and trajectory files and for detection of hydrogen bonds (hydrogen bond analysis module). The most computing-intensive part, distance calculation for the detection of hydrophobic contacts, adopts a parallel implementation with Numba (https://numba.pydata.org/) and can be accelerated on multiple CPU cores. The code is available at https://github.com/reuter-group/MD-contacts-analysis. The analysis of protein (apo and holo) in water trajectories was performed on the entire duration of the trajectories. However, for the protein-membrane simulation systems, only the last 500 ns were utilized for analysis, except for the time-series analysis. The root-mean-square deviation analysis was done on backbone atoms of protein with respect to the X-ray structure. Residues within the range of 105 to 111 were designated as Ω1 loop for the radius of gyration analysis. A contact is defined when any atom of the Ω1 loop is within a distance less than or equal to 4 Å from any atom of the C-terminal helix (residues 186-205). The criteria for hydrogen bonds are as follows: the distance between the hydrogen atom and the acceptor atom should be equal to or less than 2.4 Å, and the angle formed by the hydrogen bond acceptor, the hydrogen and hydrogen bond donor should be equal to or greater than 130 degrees. Additionally, these criteria for hydrogen bond must be met for at least two consecutive frames and present in at least two replicas. The depth of insertion was determined by analyzing the last 500 ns of the simulation trajectories. For each amino acid, the depth of insertion is calculated as the average distance between its Cα atom and the average upper phosphate plane. Cation-π interactions between the aromatic rings of aromatic residues and choline head groups were deemed to occur when the distances between the aromatic carbon and the choline nitrogen were all below 7 Å. Furthermore, these distances should not differ by more than 1.5 Å. Hydrophobic contacts are deemed to exist when two non-bonded candidate atoms are within a distance of 3 Å or less for a minimum of two consecutive frames and present in all replicas (candidate atoms for hydrophobic contacts are listed in Table S2). The tilt angle was defined as the angle between the membrane normal and the axis of helix α4, which was itself defined as a vector connecting the Cα atoms of Asp187 and Asp200. We generated data visualizations using Matplotlib, a widely-used 2D graphics library for Python(66). All graphical representations were rendered using VMD software (version 1.9.3)(37).

### Liposome micro-array based assay (LiMA)

#### LTP expression and cell extract preparation

STARD4 (RefSeq NM_139164.1) was expressed as an N-terminal His6-SUMO3-sfGFP fusion in *E. coli* (BL21 STAR; Invitrogen)(67). Expression and solubility of the fusion proteins were assessed using western blot. Cell lysis was performed as described previously(68).

#### LiMA experimental procedure

The fabrication of the liposome microarrays, the protein-liposome interaction assay, the image analysis, and the calculation of normalized binding intensity (NBI) values were performed as described previously(68). In brief, liposomes were formed in the assay buffer (10 mM HEPES (pH 7.4), 150 mM NaCl) from lipid mixtures containing 5 or 10 mol% DOPI(3)P, PI(4)P brain, DOPI(5)P, DOPI(3,4)P_2_, DOPI(3,5)P_2_, DOPI(4,5)P2, PI(4,5)P2 brain, and DOPI(3,4,5)P_3_ in a DOPC, POPC or PM-mimic background (outer PM: 33 mol% of POPC, SM, and cholesterol; inner PM: 32 mol% POPC, 12 mol% POPE, 37 mol% cholesterol, and 19 mol% POPS). Lipids were purchased form Avanti Polar Lipids and Sigma (cholesterol). The liposomes were incubated with cell extracts containing sfGFP-tagged STARD4. Subsequently, the unbound material was washed out, and the interactions were monitored by automated microscopy. Fluorescence intensities from pixels matching liposomal membranes were extracted and used for calculating the NBIs.

## Data analysis

The NBI values were divided by the NBI value of the corresponding PC (DOPC or POPC) spot on the same chip to account for chip-to-chip variability and log-transformed. Data was plotted as mean ± SEM and the individual data points for each condition. An unpaired t-test with Welch’s correction or a Brown-Forsythe ANOVA followed by a Dunnett’s t-test for multiple comparisons was performed on the data in Graphpad Prism (v10.0.2).

## Supporting information

This article contains supporting information.

## Data availability

All the MD trajectories are uploaded to the Norwegian national infrastructure for research data (NIRD), have been issued a DOI (10.11582/2023.00138) and can be accessed using the following URL: https://doi.org/10.11582/2023.00138

## Declaration of interests

The authors declare that they have no conflicts of interest with the contents of this article.

## Funding and additional information

NR and RT acknowledge funding from the Research Council of Norway (Norges Forskningsråd, grants #288008 and #335772). Computational resources were provided by NRIS/Sigma2 (#nn4700k). ACG and LvE acknowledges the financial support of Louis-Jeantet Foundation.

### Abbreviations

STARD4: Steroidogenic Acute Regulatory Protein-related Lipid-Transfer Domain 4
PIP2: Phosphatidylinositol biphosphates
LTPs: Lipid transfer proteins
StART: Steroidogenic acute regulatory protein (StAR)-related lipid-transfer
PM: Plasma membrane
ER: Endoplasmic reticulum
PI(4,5)P2: Phosphatidylinositol 4,5-bisphosphate
HMMM: highly mobile membrane-mimetic
POPC: 1-palmitoyl-2-oleoyl-sn-glycero-3-phosphocholine
POPE: 1-palmitoyl-2-oleoyl-sn-glycero-3-phosphoethanolamine
POPS: 1-palmitoyl-2-oleoyl-sn-glycero-3-phospho-L-serine
PSM: N-Palmitoylsphingomyelin
Chol: Cholesterol
POPI: 1-palmitoyl-2-oleoyl-sn-glycero-3-phosphoinositol
CER180: N-stearoyl-D-erythro-sphingosine
DOPC: 1,2-dioleoyl-sn-glycero-3-phosphocholine
DOPE: 1,2-dioleoyl-sn-glycero-3-phosphoethanolamine
DOPS: 1,2-dioleoyl-sn-glycero-3-phospho-L-serine
PLPC: 1-palmitoyl-2-linoleoyl-sn-glycero-3-phosphocholine
SAPC: 1-stearoyl-2-arachidonoyl-sn-glycero-3-phosphocholine
PSPE: 1-Stearoyl-2-palmitoyl-sn-glycero-3-phosphoethanolamine
SAPE: 1-stearoyl-2-arachidonoyl-sn-glycero-3-phosphoethanolamine
SAPI: 1-stearoyl-2-arachidonoyl-sn-glycero-3-phosphoinositol
SLPI: 1-Stearoyl-2-linoleoyl-sn-glycero-3-phosphoinositol
POPA: 1-palmitoyl-2-oleoyl-sn-glycero-3-phosphate
LiMA: Liposome micro-array based assay
NBI: Normalized binding intensity

## Notes

### Competing Interest Statement

The authors have declared no competing interest.

