## Supporting Information for "Membrane specificity of the human cholesterol transfer protein STARD4"

[1] Department of Chemistry, University of Bergen, Bergen, 5020, Norway
[2] Computational Biology Unit, Department of Informatics, University of Bergen, Bergen, 5020, Norway
[3] Department of Cellular Physiology and Metabolism, Faculty of Medicine, University of Geneva, Geneva Switzerland
[4] Diabetes Center, Faculty of Medicine, University of Geneva, Geneva, Switzerland
[5] École Polytechnique Fédérale de Lausanne EPFL, Lausanne, Switzerland

**Corresponding author*


### Material included:

**Table S1:** List of simulated systems.

**Table S2:** List of atoms considered for analysis of hydrophobic contacts.

**Figure S1:** STARD4 structure, topology and sequence.

**Figure S2:** Secondary structure analysis of C-ter helix.

**Figure S3:** Root mean square deviation (RMSD) analysis.

**Figure S4:** Minimum distance between protein and lipid bilayers along simulation time.

**Figure S5:** Depth of insertion plot for each amino acid in STARD4.

**Figure S6:** STARD4-PI(4,5)P2 hydrogen bonds.

**Figure S7:** Lipid snorkeling in STARD4-bilayers simulations.

**Movie S1:** Movie of STARD4 simulated on the plasma membrane bilayer model.

**Table S1**. List of simulated systems. Each system was simulated for 2 μs (in addition to the equilibration steps).

| Simulation type | Protein state | Lipid bilayer | Acronym | No. of Replicas | No. of Water | Total No. of Atoms | No. of Ions |
|---|---|---|---|---|---|---|---|
| Water | Apo | - | Apo-W | Rep-1 | 11759 | 38479 | 5 $Na^+$ |
| | | | | Rep-2 | 13869 | 44809 | |
| | Holo | - | Holo-W | Rep-1 | 13799 | 44673 | 5 $Na^+$ |
| | | | | Rep-2 | 13348 | 43320 | |
| Plasma membrane | Apo | nPM | Apo-nPM | Rep-1 | 41949 | 170409 | 57 $Na^+$ |
| | | | | Rep-2 | 37196 | 156150 | |
| | Apo | mPM | Apo-mPM | Rep-1 | 40512 | 165942 | 39 $Na^+$ |
| | | | | Rep-2 | 36537 | 154017 | |
| | Holo | nPM | Holo-nPM | Rep-1 | 39418 | 162890 | 57 $Na^+$ |
| | | | | Rep-2 | 39401 | 162839 | |
| Endoplasmic reticulum (ER) | Apo | pER | Apo-pER | Rep-1 | 31583 | 130785 | 45 $Na^+$ |
| | | | | Rep-2 | 31580 | 130776 | |
| | Holo | pER | Holo-pER | Rep-1 | 30965 | 129005 | 45 $Na^+$ |
| | | | | Rep-2 | 30924 | 128882 | |
| | | | | Rep-3 | 30481 | 127553 | |
| | | | | Rep-4 | 29641 | 125033 | |
| | Holo | dER | Holo-dER | Rep-1 | 30806 | 129360 | 45 $Na^+$ |
| | | | | Rep-2 | 29157 | 124413 | |
| | Holo | cER | Holo-cER | Rep-1 | 29895 | 126545 | 33 $Na^+$ |
| | | | | Rep-2 | 28173 | 121379 | |
| | Holo | hER | Holo-hER | Rep-1 | 32893 | 129253 | 45 $Na^+$ |
| | | | | Rep-2 | 31122 | 125432 | |
| Pure POPC | Apo | PC | Apo-PC | Rep-1 | 35672 | 144522 | 5 $Na^+$ |
| | | | | Rep-2 | 33439 | 137897 | |

**Table S2**. **List of atoms considered for analysis of hydrophobic contacts.** The atom names correspond to the nomenclature of the CHARMM36m force field.

| Residue/lipid | Candidate atoms |
|---|---|
| ALA | CA HA CB HB1 HB2 HB3 |
| ARG | CA HA CB HB1 HB2 CG HG1 HG2 |
| ASN | CA HA CB HB1 HB2 |
| ASP | CA HA CB HB1 HB2 |
| CYS | CA HA CB HB1 HB2 |
| GLN | CA HA CB HB1 HB2 CG HG1 HG2 |
| GLU | CA HA CB HB1 HB2 CG HG1 HG2 |
| GLY | CA HA1 HA2 |
| HSD | CA HA CB HB1 HB2 |
| ILE | CA HA CB HB CG1 HG11 HG12 CG2 HG21 HG22 HG23 CD HD1 HD2 HD3 |
| LEU | CA HA CB HB1 HB2 CG HG CD1 HD11 HD12 HD13 CD2 HD21 HD22 HD23 |
| LYS | CA HA CB HB1 HB2 CG HG1 HG2 CD HD1 HD2 CE HE1 HE2 |
| MET | CA HA CB HB1 HB2 CG HG1 HG2 CE HE1 HE2 HE3 |
| PHE | CA HA CB HB1 HB2 CG CD1 HD1 CD2 HD2 CE1 HE1 CE2 HE2 CZ HZ |
| PRO | CA HA CB HB1 HB2 CD HD1 HD2 CG HG1 HG2 |
| SER | CA HA CB HB1 HB2 |
| THR | CA HA CB HB CG2 HG21 HG22 HG23 |
| TRP | CA HA CB HB1 HB2 CG CD1 HD1 CD2 CE3 HE3 CZ3 HZ3 CH2 HH2 CZ2 HZ2 |
| TYR | CA HA CB HB1 HB2 CG CD1 HD1 CD2 HD2 CE1 HE1 |
| VAL | CA HA CB HB CG1 HG11 HG12 HG13 CG2 HG21 HG22 HG23 |
| POPC/POPE/ POPS/POPI/ POPA | C23 H3R H3S C24 H4R H4S C25 H5R H5S C26 H6R H6S C27 H7R H7S C28 H8R H8S C29 H91 C210 H101 C211 H11R H11S C212 H12R H12S C213 H13R H13S C214 H14R H14S C215 H15R H15S C216 H16R H16S C217 H17R H17S C218 H18R H18S H18T C33 H3X H3Y C34 H4X H4Y C35 H5X H5Y C36 H6X H6Y C37 H7X H7Y C38 H8X H8Y C39 H9X H9Y C310 H10X H10Y C311 H11X H11Y C312 H12X H12Y C313 H13X H13Y C314 H14X H14Y C315 H15X H15Y C316 H16X H16Y H16Z |
| PLPC | C23 H3R H3S C24 H4R H4S C25 H5R H5S C26 H6R H6S C27 H7R H7S C28 H8R H8S C29 H9S C21S H10S C211 H11R H11S C212 H12R C213 H13R C214 H14R H14S C215 H15R H15S C216 H16R H16S C217 H17R H17S C218 H18R H18S H18T C33 H3X H3Y C34 H4X H4Y C35 H5X H5Y C36 H6X H6Y C37 H7X H7Y C38 H8X H8Y C39 H9X H9Y C310 H10X H10Y C311 H11X H11Y C312 H12X H12Y C313 H13X H13Y C314 H14X H14Y C315 H15X H15Y C316 H16X H16Y H16Z |
| PSPE | C23 H3R H3S C24 H4R H4S C25 H5R H5S C26 H6R H6S C27 H7R H7S C28 H8R H8S C29 H9S H9R C210 H10S H10R C211 H11R H11S C212 H12R H12S C213 H13R H13S C214 H14R H14S C215 H15R H15S C216 H16R H16S C217 H17R H17S C218 H18R H18S H18T C33 H3X H3Y C34 H4X H4Y C35 H5X H5Y C36 H6X H6Y C37 H7X H7Y C38 H8X H8Y C39 H9X H9Y C310 H10X H10Y C311 H11X H11Y C312 H12X H12Y C313 H13X H13Y C314 H14X H14Y C315 H15X H15Y C316 H16X H16Y H16Z |

| | |
|---|---|
| DOPC/DOPE/ DOPS | C12 C11 H11A H11B C1 HA HB C2 HS C22 H2R H2S C3 HX HY C32 H2X H2Y C23 H3R H3S C24 H4R H4S C25 H5R H5S C26 H6R H6S C27 H7R H7S C28 H8R H8S C29 H9R C210 H10R C211 H11R H11S C212 H12R H12S C213 H13R H13S C214 H14R H14S C215 H15R H15S C216 H16R H16S C217 H17R H17S C218 H18R H18S H18T C33 H3X H3Y C34 H4X H4Y C35 H5X H5Y C36 H6X H6Y C37 H7X H7Y C38 H8X H8Y C39 H9X C310 H10X C311 H11X H11Y C312 H12X H12Y C313 H13X H13Y C314 H14X H14Y C315 H15X H15Y C316 H16X H16Y C317 H17X H17Y C318 H18X H18Y H18Z |
| SAPC/SAPE/ SAPI/SAPI24 | C11 H1 C12 H2 C13 H3 C14 H4 C15 H5 C16 H6 C1 HA HB C2 H3 C3 HX HY C22 H2S H2R C23 H3S H3R C24 H4S H4R C25 H5R C26 H6R C27 H7S H7R C28 H8R C29 H9R C210 H10S H10R C211 H11R C212 H12R C213 H13S H13R C214 H14R C215 H15R C216 H16S H16R C217 H17S H17R C218 H18S H18R C219 H19S H19R C220 H20S H20R H20T C32 H2X H2Y C33 H3X H3Y C34 H4X H4Y C35 H5X H5Y C36 H6X H6Y C37 H7X H7Y C38 H8X H8Y C39 H9X H9Y C310 H10X H10Y C311 H11X H11Y C312 H12X H12Y C313 H13X H13Y C314 H14X H14Y C315 H15X H15Y C316 H16X H16Y C317 H17X H17Y C318 H18X H18Y H18Z |
| SLPI | C12 C11 H11A H11B C1 HA HB C2 HS C22 H2R H2S C3 HX HY C32 H2X H2Y C23 H3R H3S C24 H4R H4S C25 H5R H5S C26 H6R H6S C27 H7R H7S C28 H8R H8S C29 H9R C210 H10R C211 H11R H11S C212 H12R C213 H13R C214 H14R H14S C215 H15R H15S C216 H16R H16S C217 H17R H17S C218 H18R H18S H18T C33 H3X H3Y C34 H4X H4Y C35 H5X H5Y C36 H6X H6Y C37 H7X H7Y C38 H8X H8Y C39 H9X H9Y C310 H10X H10Y C311 H11X H11Y C312 H12X H12Y C313 H13X H13Y C314 H14X H14Y C315 H15X H15Y C316 H16X H16Y C317 H17X H17Y C318 H18X H18Y H18Z |
| OLPS | C12 C11 H11A H11B C1 HA HB C2 HS C22 H2R H2S C3 HX HY C32 H2X H2Y C23 H3R H3S C24 H4R H4S C25 H5R H5S C26 H6R H6S C27 H7R H7S C28 H8R H8S C29 H9R C210 H10R C211 H11R H11S C212 H12R C213 H13R C214 H14R H14S C215 H15R H15S C216 H16R H16S C217 H17R H17S C218 H18R H18S H18T C33 H3X H3Y C34 H4X H4Y C35 H5X H5Y C36 H6X H6Y C37 H7X H7Y C38 H8X H8Y C39 H9X C310 H10X C311 H11X H11Y C312 H12X H12Y C313 H13X H13Y C314 H14X H14Y C315 H15X H15Y C316 H16X H16Y C317 H17X H17Y C318 H18X H18Y H18Z |
| CER180 | C5S C6S H6S H6T C7S H7S H7T C8S H8S H8T C9S H9S H9T C10S H10S H10T C11S H11S H11T C12S H12S H12T C13S H13S H13T C14S H14S H14T C15S H15S H15T C16S H16S H16T C17S H17S H17T C18S H18S H18T H18U C2F H2F H2G C3F H3F H3G C4F H4F H4G C5F H5F H5G C6F H6F H6G C7F H7F H7G C8F H8F H8G C9F H9F H9G C10F H10F H10G C11F H11F H11G C12F H12F H12G C13F H13F H13G C14F H14F H14G C15F H15F H15G C16F H16F H16G C17F H17F H17G C18F H18F H18G H18H |
| PSM | C11 H11A H11B C12 H12A H12B C13 H13A H13B H13C C14 H14A H14B H14C C15 H15A H15B H15C C1S H1S H1T C2S H2S C3 H3S C2F H2F H2G C3F H3F H3G C4F H4F H4G C5F H5F H5G C6F H6F H6G C7F H7F H7G C8F H8F H8G C9F H9F H9G C10F H10F H10G C11F H11F H11G C12F H12F H12G C13F H13F H13G C14F H14F H14G C15F H15F H15G C16F H16F H16G H16H C4S H4S C5S H5S C6S H6S H6T C7S H7S H7T C8S H8S H8T C9S H9S H9T C10S H10S H10T C11S H11S H11T C12S H12S H12T C13S H13S H13T C14S H14S |

| | |
|---|---|
| | H14T C15S H15S H15T C16S H16S H16T C17S H17S H17T C18S H18S H18T H18U |
| Cholesterol | C4 H4A H4B C5 C6 H6 C7 H7A H7B C8 H8 C14 H14 C15 H15A H15B C16 H16A H16B C17 H17 C13 C18 H18A H18B H18C C12 H12A H12B C11 H11A H11B C9 H9 C10 C19 H19A H19B H19C C1 H1A H1B C2 H2A H2B C20 H20 C21 H21A H21B H21C C22 H22A H22B C23 H23A H23B C24 H24A H24B C25 H25 C26 H26A H26B H26C C27 H27A H27B H27C |

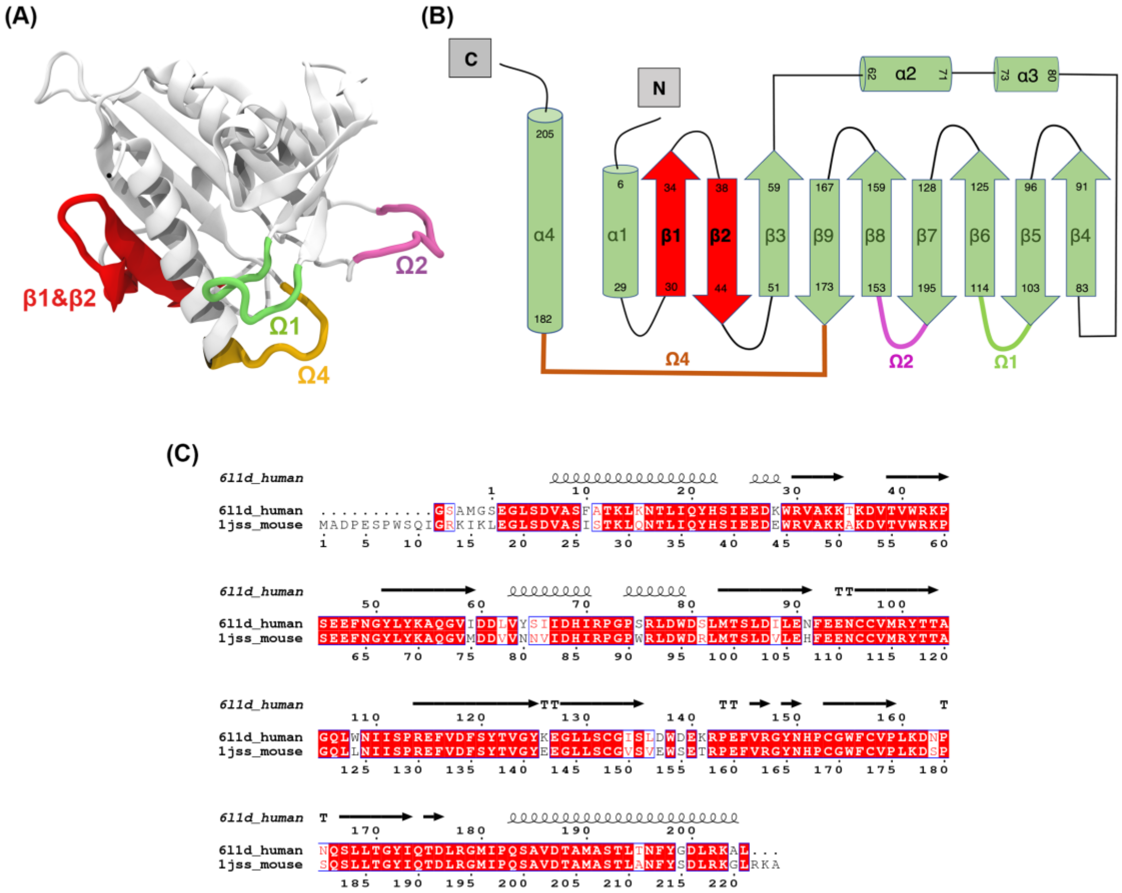


**Figure S1. STARD4 structure, topology and sequence. (A)** Tertiary structure of STARD4 in cartoon model. **(B)** Secondary structure diagram **(C)** Sequence alignment of human and mouse STARD4.

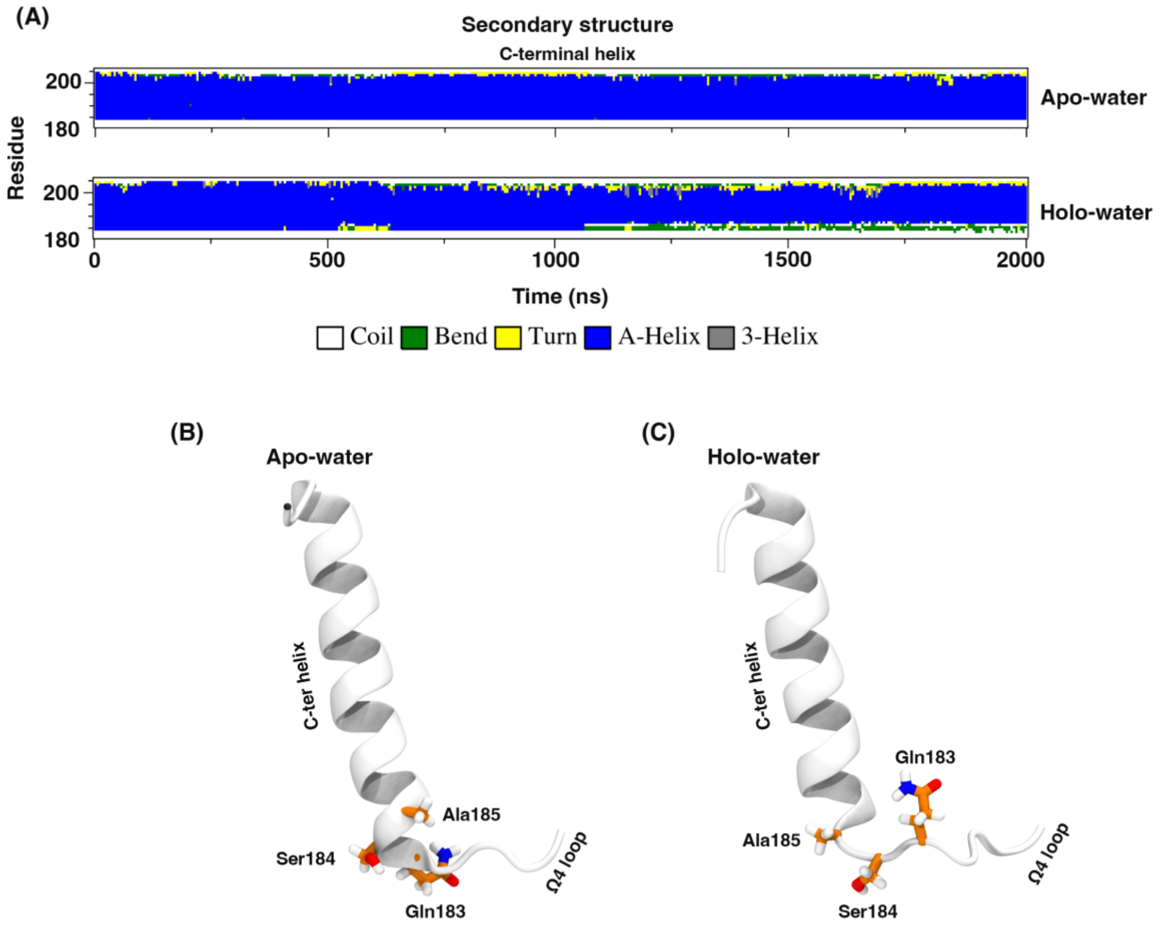


**Figure S2. Secondary structure analysis of C-ter helix in simulations of STARD4 in water. (A)** Secondary structure analysis of the C-terminal helix and Ω4 loop of the apo and holo protein structures. **(B-C)** Structure of the C-terminal helix at the end of the apo-water (B) and of the holo-water (C) simulations. The C-terminal helix and the Ω4 loop are represented using cartoons (white). The carbon, oxygen, nitrogen, and hydrogen atoms of amino acids 183-185 are colored orange, red, blue, and white respectively.

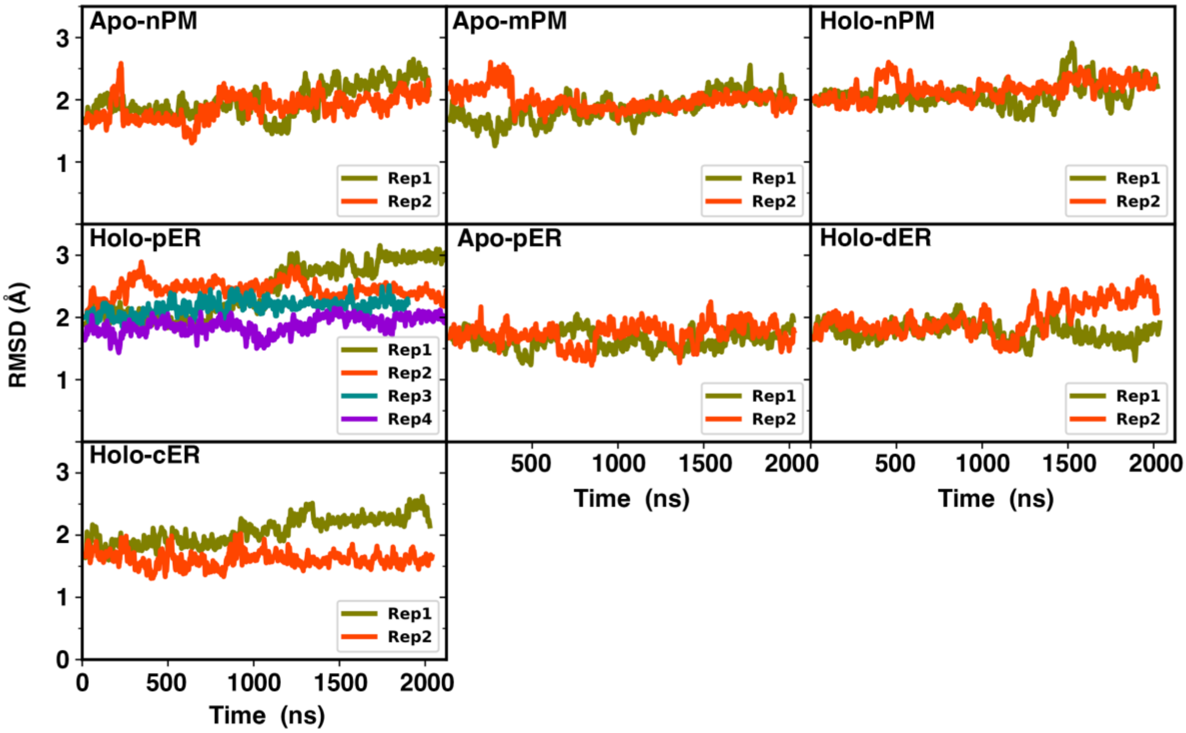


**Figure S3.** Root mean square deviation (RMSD) of protein backbone atoms with respect to the X-ray structure in all protein-membrane simulation systems.

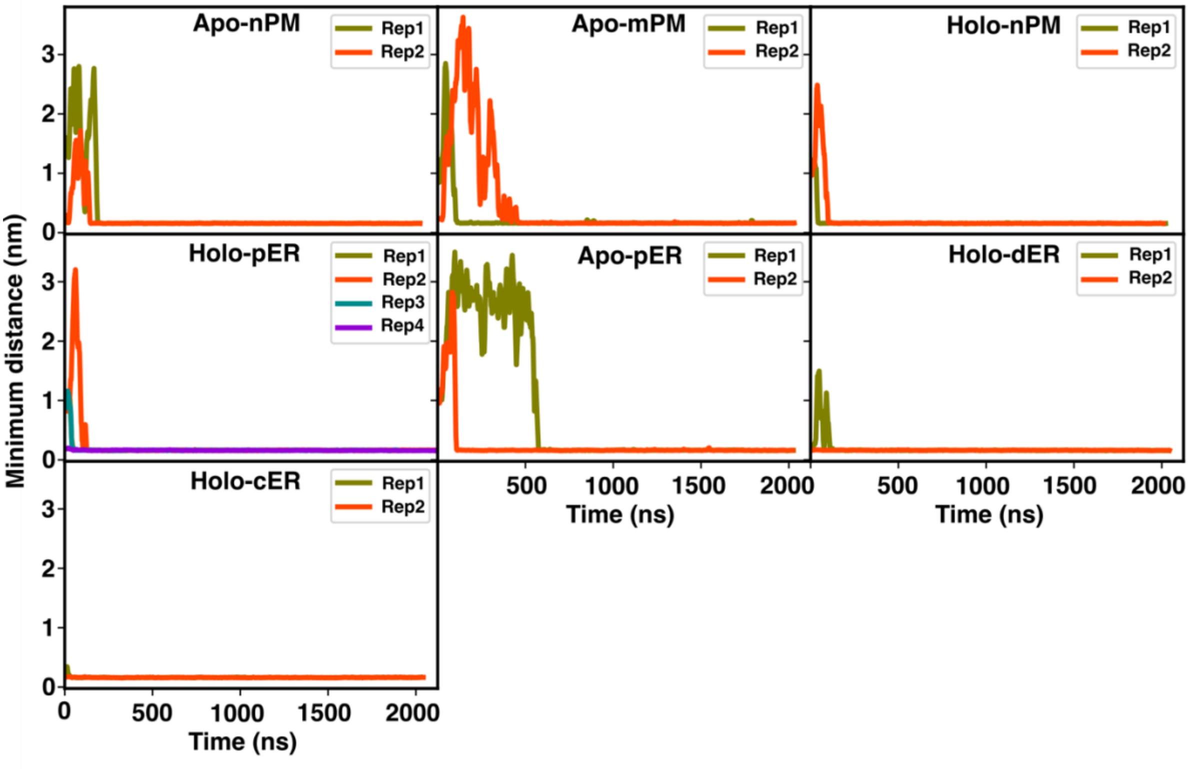


**Figure S4.** Minimum distance between protein and lipid bilayers along simulation time.

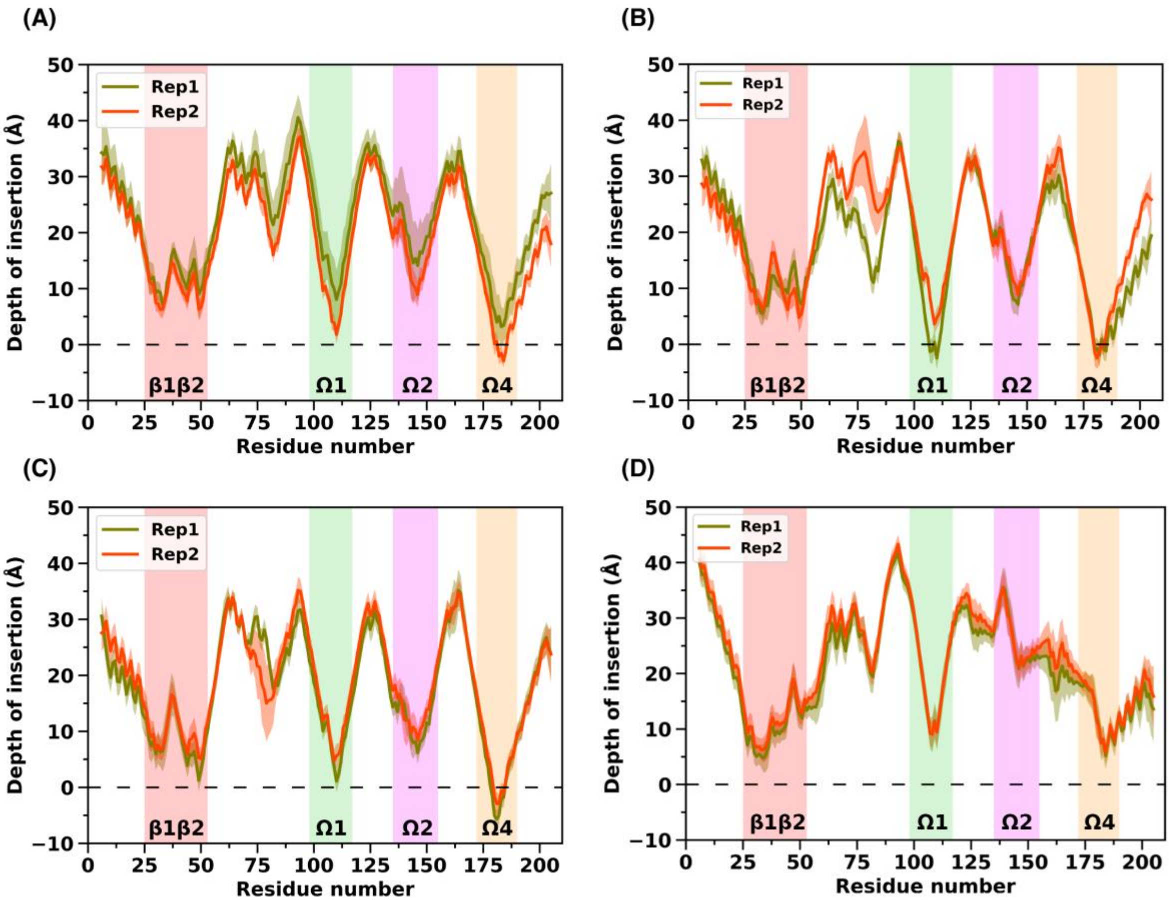


**Figure S5.** Depth of insertion plot for each amino acid in STARD4. **(A)** Holo-nPM, **(B)** Holo-dER, **(C)** Holo-cER, and **(D)** Apo-PC simulation systems. The data are averaged over the last 500 ns of each simulation.

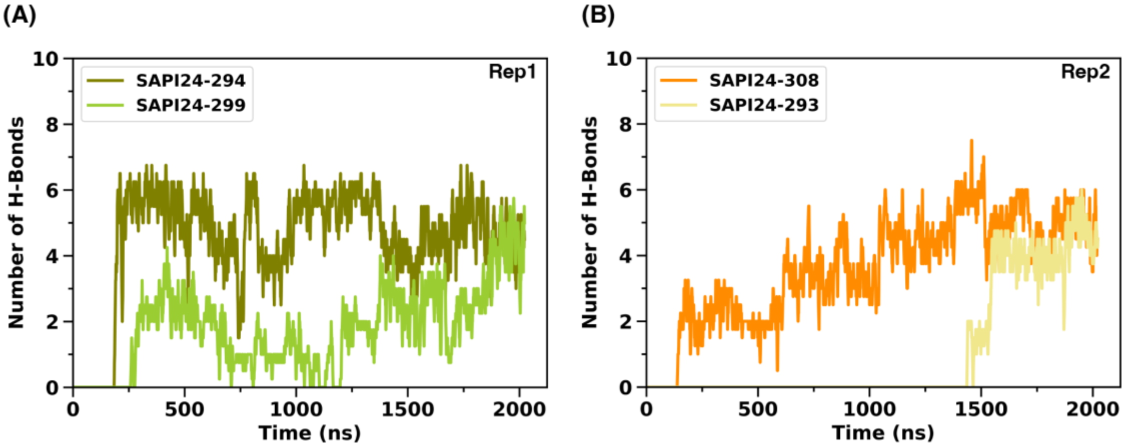


**Figure S6. STARD4-PIP2 hydrogen bonds. (A-B)** number of hydrogen bonds between protein residues and selected PI(4,5)P2 lipids in the Apo-nPM system along two replica trajectories.

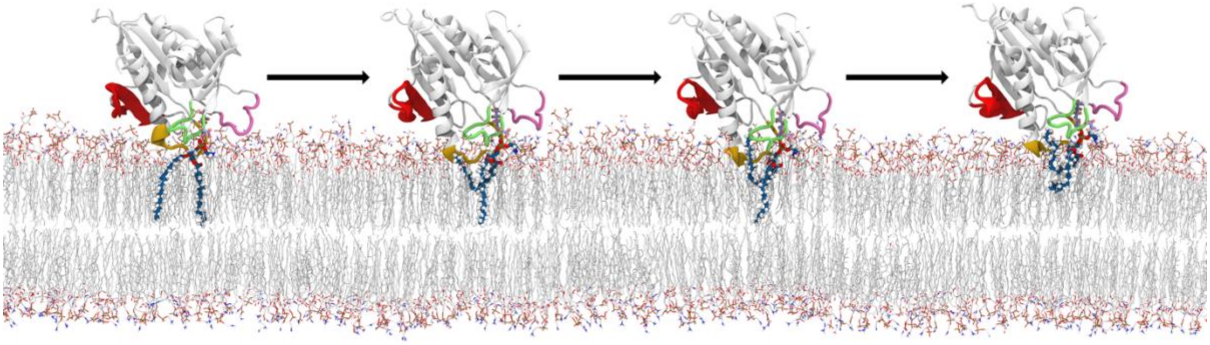

**Figure S7. Lipid snorkeling in STARD4-bilayers simulations.** The lipid interacting with Ser112 and Arg114 (through its polar head) snorkels and inserts its unsaturated tail in the gap between the Ω1 and Ω4 loops.

**Movie S1. Movie of STARD4 simulated on the plasma membrane bilayer model (Apo-nPM) (0 to 250 ns).** The protein is shown and colored as on Figure 5, basic residues in the PIP2 binding sites are represented as licorice (orchid violet), nPM bilayer lipids are shown as lines (carbon, nitrogen, phosphorous, and oxygen atoms are colored white, blue, tan, and red, respectively). Two PI(4,5)P2 lipids are highlighted using licorice (carbon, phosphorous, and oxygen atoms are colored orange, tan, and red, respectively). Hydrogen atoms are not shown for the sake of clarity.
